# Etiology of Craniofacial and Cardiac Malformations in a Mouse Model of SF3B4-Related Syndromes

**DOI:** 10.1101/2024.01.30.578039

**Authors:** Shruti Kumar, Eric Bareke, Jimmy Lee, Emma Carlson, Fjodor Merkuri, Evelyn E. Schwager, Steven Maglio, Jennifer L. Fish, Jacek Majewski, Loydie Jerome-Majewska

## Abstract

Pathogenic variants in SF3B4 are responsible for the acrofacial disorders Nager and Rodriguez Syndrome, also known as SF3B4-related Syndromes, associated with malformations in the head, face, limbs, vertebrae as well as the heart. To uncover the etiology of craniofacial malformations found in SF3B4-related syndromes, mutant mouse lines with homozygous deletion of *Sf3b4* in neural crest cells (NCC) were generated. Like in human patients, these embryos had craniofacial and cardiac malformations with variable expressivity and penetrance. The severity and survival of *Sf3b4* NCC mutants was modified by the level of *Sf3b4* in neighbouring non NCC. RNA sequencing analysis of heads of embryos prior to morphological abnormalities showed significant changes in expression of genes forming the NCC regulatory network, as well as an increase in exon skipping. We identified several key transcription factors and histone modifiers involved in craniofacial and cardiac development with increased exon skipping. Increased exon skipping was also associated with use of a more proximal branch point, as well as an enrichment in thymidine bases in the 50bp around the branch points. We propose that decrease in *Sf3b4* causes changes in the expression and splicing of transcripts required for proper craniofacial and cardiac development, leading to abnormalities.

## INTRODUCTION

In humans, pathogenic variants in *SF3B4* are responsible for the most common type of acrofacial dysostosis, Nager syndrome^1,2^ (OMIM #154400) and Rodriguez syndrome^3^ (OMIM#201170), here on referred to as SF3B4-related disorders^4^. Acrofacial Dysostosis and SF3B4-related disorders are characterized by micrognathia and ear defects as well as limb defects. Craniofacial and pre-axial limb defects common in SF3B4 patients include; malar and mandibular hypoplasia, cleft palate, downward slanted palpebral fissures, hearing loss, small or absent thumbs, and minor foot anomalies such as small metatarsals^1,3,5,6^. Additionally, some patients exhibit rib abnormalities and abnormal vertebral segmentation^1,3^, as well as ventricular septal defect, coarctation of the aorta, hypertrabeculation of the left ventricle and complex structural heart disease^1,5,6^. 41 distinct pathogenic variants have been identified in *SF3B4* to date^1–14^, and include whole gene deletion, nonsense, missense, frameshifts, and truncating changes. Since no genotype-phenotype correlation has been found^15^, all mutations are predicted to be non-functional.

The splicing factor *SF3B4* encodes for splicing associated protein (SAP49)/SF3B4 and is a core component of the SF3B complex found in both the U2 as well as the U12 small nuclear ribonucleoproteins (snRNP)^16^. SF3B4 has two RNA recognition motifs (RRM) at its amino terminal that facilitates binding of U2 immediately upstream of the 3’ branch point adenosine during spliceosome assembly^17^. The major or U2-dependent spliceosome catalyses 99% of the RNA splicing reactions in humans^18^, whereas the U12 dependent spliceosome (minor spliceosome) is responsible for splicing of 700 minor introns in 666 genes^19^. The spliceosome is also crucial for alternative splicing, an important contributor to protein and genetic diversity^16,20^, that is indispensable for development and maintenance of complex organisms.

Our aim is to uncover the etiology of malformations seen in SF3B4-related syndromes. Previous work by our group and the group of Yamada et al^21,22^, showed that mice with heterozygous mutation of *Sf3b4* did not model craniofacial malformations seen in patients, but had growth retardation, microcephaly, as well as homeotic posteriorization of the axial skeleton. Furthermore, although craniofacial and eye abnormalities found in SF3B4*-*related patients have been phenocopied in zebrafish and *Xenopus* embryos with knock down of *Sf3b4*^23,24^, vertebrae and heart defects were not found. Additionally, though RNAseq analysis showed few changes in gene expression, a significant number of genes including histone modifiers that regulate *Hox* expression, were abnormally spliced in heterozygous mouse embryos and somites^21^.

Since structures affected in the face and head of patients are all derived from cranial neural crest cells (CNCC), herein, we generated a mouse model with conditional mutation in *Sf3b4* and used the *Wnt1-Cre2* line to delete *Sf3b4* in neural crest cells (NCC). *Sf3b4* NCC mutant embryos exhibit craniofacial, heart and vertebral abnormalities similar to those described in patients with SF3B4*-*related diseases. These malformations were associated with decreased expression of NCC specifier genes, as well as aberrant splicing of histone modifiers important for craniofacial development. For the first time, we identified sequence specific changes associated with abnormal branch point recognition and mis-splicing in *Sf3b4* mutants. Overall, we show that *Sf3b4* is necessary at the earliest stages of NCC differentiation and propose that transcripts with branch-points situated in thymine-rich contexts have an increased susceptibility to be mis-spliced when levels of SF3B4 are reduced.

## RESULTS

### Deletion of *Sf3b4* with *Wnt1-Cre2* results in craniofacial malformations that are modified by the genotype of neighboring cells

Previously generated *Sf3b4* heterozygous mutant embryos on two genetic backgrounds showed vertebrae abnormalities but did not have craniofacial defects^21,22^. Here, to characterize the requirement for *Sf3b4* during craniofacial development and the etiology of craniofacial malformations in SF3B4*–*related syndromes, *Sf3b4* conditional mutant mice with loxP sequences flanking exons 2 – 3 of *Sf3b4 (Sf3b4^L/+^*)^21^ and *Wnt1-Cre*2^25^ were used to delete *Sf3b4* in NCC (Figure S1A and S1B). The *Wnt1* promoter drives CRE expression and activity in the neuroepithelium and NCC that emerges from the forebrain and midbrain of E8.5 embryos with 4 or more somites (s)^25,26^.

In previously generated mice with NCC mutation in the splicing genes *Eftud2* and *Snrpb*, craniofacial defects are first seen at E9.5 and few mutants survive to birth^27,28^. To determine when *Sf3b4* NCC mutants are first distinguishable from their control littermates, embryos generated from matings between *Sf3b4^L/+^* and *Sf3b4^L/+^;Wnt^tg/+^* (giving *Sf3b4^ncc/ncc^* mutant embryos: *Sf3b4^L/L^ ;Wnt^tg/+^*), or *Sf3b4^+/-^;Wnt^tg/+^* (giving *Sf3b4^ncc/–^* mutant embryos: *Sf3b4^L/-^ ;Wnt^tg/+^*) mice were analyzed from embryonic day (E) 8.5 – E17.5. At E8.5, only 72% (n=34/47) of *Sf3b4^ncc/ncc^* embryos and 53% (n= 55/103) of *Sf3b4^ncc/–^* mutants were normal, compared to 99% (n=366/370) of somite-matched controls (*Sf3b4^+/+^;Wnt1-Cre2^tg/+^*, *Sf3b4^+/–^*, and *Sf3b4^ncc/+^*). Though hypoplasia of the first pharyngeal arch and the fore- and mid-brain were the most common abnormalities found in mutants, as observed in 96% (n=46/48) of *Sf3b4^ncc/–^* and 54% (n=7/13) of *Sf3b4^ncc/ncc^* mutant embryos, phenotypic differences were found in these two genotypes. Specifically, 8% (n=4/48) of *Sf3b4^ncc/–^* mutants exhibited hypoplasia of the heart, a phenotype not previously seen in spliceosomal mouse mutants, and *Sf3b4^ncc/ncc^* mutants exhibited either hypoplasia of only the 1^st^ pharyngeal arch (23% (n=3/13), or hypoplasia of only the midbrain (23% (n=3/13). A day later, at E9.5 the majority of *Sf3b4* NCC mutants for both genotypes were abnormal (Sup Table 1.1 and 1.2) and although 2% (n=1/45) of *Sf3b4^ncc/–^* mutants were undergoing resorption, all E9.5 *Sf3b4^ncc/ncc^* mutants (n=22) were alive. Thus, *Sf3b4* NCC mutants can first be distinguished from their control littermates at E8.5, shortly after CRE expression.

**Table 1:** Transcripts with differentially skipped exons that have gene ontology of histone modifier and have craniofacial/heart phenotype.

| Gene | Skipped Exon | Function | Phenotype | Reference |
| --- | --- | --- | --- | --- |
| *Tcf7l2* | 4 | Transcription Factor | Abnormal brain and craniofacial development | Jan Masek et al, 2016<br>doi:10.1242/dev.132357 |
| *Kdm5c* | 24 | H3K4 demethylase | Small head, reduced cartilage, eye abnormality | Kim et al, 2018<br>https://doi.org/10.1186/s13072-018-0241-x |
| *Kdm6a* | 11 | H3K27 demethylase | Cleft palate, abnormal heart and brain development | Shpargel et al., 2017<br>https://doi.org/10.1073/pnas.1705011111 |
| *Setd5* | 5 | H3K36 trimethylase | Small head, brain and eye abnormality | Osipovich et al, 2016<br>doi:10.1242/dev.141465 |
| *Carm1* | 15 | Arginine methyltransferase | Smaller, delayed ossification | Ito et al, 2009<br>doi:10.1186/1471-213X-9-47 |
| *Dnmt3a* | 9 | Cytosine methyltransferase | Not required for embryonic development but required for neural crest cell development | Hu et al, 2012<br>doi: 10.1101/gad.198747.112 |
| *Dnmt3b* | 4 | Cytosine methyltransferase | Abnormal craniofacial morphology, cleft palate | Ueda et al, 2006<br>https://doi.org/10.1242/dev.02293 |

From E10.5 to E17.5, *Sf3b4* NCC mutants (n= 74, *Sf3b4^ncc/ncc^* and 62 *Sf3b4^ncc/–^*) were assessed at the time of dissection for presence of a heartbeat and phenotypic defects. Most *Sf3b4* NCC mutants were alive, as only 21% (n=28/136) were undergoing resorption. Live mutant embryos exhibited a spectrum of defects which we grouped into three phenotypic classes (Figure S1C): Class 1 - appeared normal; Class 2 (moderate), exhibited hypoplasia of the developing pharyngeal arches, or of the brain, or an open neural tube; and Class 3 (severe). Class 3 embryos found between E10.5 – E12.5 exhibited hypoplasia of both the brain and pharyngeal arches, or reduced frontonasal mass, and/or mandible, while E13.5 – E17.5 embryos in this class showed anencephaly – an exposed forebrain, absent eyes and ears, as well as hypoplastic mandible. Furthermore, all E17.5 *Sf3b4* NCC mutants weighed significantly less than their control litter mates (Figure S2F). From a total of 136 *Sf3b4* NCC mutants collected from E10.5-17.5, Class 1 mutants were present at all stages with 20%, being *Sf3b4^ncc/ncc^*, (n=27/136) while 13% (n=8/136) were *Sf3b4^ncc/–^*. Similarly, whereas 10% (n=14/136) of *Sf3b4* NCC mutant embryos belonging to Class 2 were *Sf3b4^ncc/ncc^* mutants, less than 1%, (1/136) were *Sf3b4^ncc/–^* mutants. In contrast, whereas 13% (18/136) of *Sf3b4* NCC Class 3 mutants were *Sf3b4^ncc/ncc^*, 29% (40/136) were *Sf3b4^ncc/–^* mutants. Thus, homozygous mutation of *Sf3b4* in NCCs, in the context of neighboring *Sf3b4* heterozygous cells, results in a more penetrant and severe phenotype.

Xenopus embryos with *Sf3b4* knockdown, showed an expanded neural tube^24^. To determine if the neural tube was also expanded in mouse E12.5 *Sf3b4* NCC mutants, the width of the left and the right neural epithelium was measured at the level of the forelimb (Figure S2A, white line). No difference was found in the width of the left and the right neural epithelium of Class 1 and 2 *Sf3b4* NCC mutants (n=4) when compared to those of controls. However, the width of both the left and the right neural folds of Class 3 *Sf3b4* NCC mutants (n=11) was significantly reduced when compared to controls (t-test, p<0.0001) (Figure S2C and S2D). Furthermore, 25% (n=1/4) of Class 1 and 2, and 64% (n=7/11) of Class 3 *Sf3b4* NCC mutants had an open neural tube (Figure S2A and SB). Thus, deletion of mouse *Sf3b4* in the dorsal neural tube results in a significant decrease in the size of the neural tube and open neural tube in a subset of embryos. When taken together, these data suggest that the reduced size of the neural epithelium may be responsible for the open neural tube found in *Sf3b4* NCC mutants.

Although the proportion of embryos undergoing resorption was similar between genotypes, 20% (n=15/74) of *Sf3b4^ncc/ncc^* and 21% (13/62) of *Sf3b4^ncc/–^*, the morphological analysis above suggests that phenotypically normal appearing embryos belonging to Class 1 may be born. To determine if this was the case, all pups found from mating of *Sf3b4^L/+^* males and *Sf3b4^L/+^;Wnt^tg/+^* females (n= 68) or *Sf3b4^L/+^* and *Sf3b4^+/–^;Wnt^tg/+^* (n=13) mice were genotyped. No *Sf3b4^ncc/–^* pups were found from these matings (Sup Table 2.2). In contrast, 2 *Sf3b4^ncc/ncc^* (Sup Table 2.1) were found at postnatal day (P) 21, when offspring were weaned. Both *Sf3b4^ncc/ncc^* female mice were indistinguishable from their wildtype litter mates and were euthanized a year after their birth. Although a significant deviation from the expected frequency, (χ^2^=17.64, p=0.0052), this data shows that mice with homozygous deletion of *Sf3b4* in NCC can be born and survive to adulthood. Additionally, these findings indicate that homozygous deletion of *Sf3b4* in NCC is not fully penetrant when all other cells of the embryo are wildtype.

To rule out a sex-specific difference in the expressivity and survival of *Sf3b4* NCC mutants to E17.5, we genotyped a subset of embryos for the sex chromosomes (n=32 *Sf3b4^ncc/ncc^* and 6 *Sf3b4^ncc/–^*). No difference was found in the proportion of male and female embryos genotyped (n=21 male and 17 female). Furthermore, no difference was found in the proportion of male and female embryos found in Class 1 (n=5 male and 7 female), Class 2 (2 male and 4 female), Class 3 (7 male and 2 female) or undergoing resorption (n=7 male and 4 female) (Sup Table 3). These data show that in mouse, like in human, the sex chromosomes do not modify expressivity and penetrance of *Sf3b4* mutation. Altogether our results show that mutation of *Sf3b4* in mouse NCC phenocopies the penetrance and expressivity of craniofacial malformations found in human patients with *SF3B4-*related syndromes.

### Neural crest and mesoderm-derived bones and cartilages are abnormally formed in *Sf3b4* NCC mutants

CNCC contribute to many of the cartilage and bones that are malformed in *SF3B4-* related syndrome patient in addition to directing differentiation of head mesoderm-derived bones and cartilage. To visualize cartilages and bones in E17.5 embryos, we used Alizarin red and Alcian blue staining. Alcian blue binds to sulfated glycosaminoglycans and glycoproteins found in cartilages, while Alizarin red binds to calcium, the main components of mineralized bones. Our analysis revealed that one Class 1 *Sf3b4^ncc/ncc^* mutant (n= 1/10) was normal and indistinguishable from its control litter mates (n=23), consistent with the fact that a subset of these mutant embryos are born and survive to adulthood. However, all remaining *Sf3b4* NCC mutants (Class 1 (n= 9), Class 2 (n=2), and Class 3 (n=7), Figure S2E) exhibited at least one defect in the craniofacial region (Sup Table 4). These defects span a broad spectrum and were variable between individual embryos of the same genotype and Class. Nonetheless, Class 3 mutant embryos had the most severe and the highest burden of defects, exhibiting multiple malformations in the skull.

Malformations found in the cranium include the absence of CNCCs and mesoderm-derived cartilage and bones ventral to the basisphenoid (n=7/7) in Class 3 embryos (Figure 1). At the other extreme, neural crest-derived structures, including the nasal cartilage, the palate, and the frontal bone were formed in the cranium of Class 1 and Class 2 mutants, but these bones had a smaller region of mineralization and were thus smaller than those of controls (Figure 1). The basisphenoid which is derived from NCC and forms via endochondral ossification was present but abnormally shaped, while the pterygoid, also NCC-derived, was absent in a subset of Class 1, Class 2 and Class 3 mutants (Figure 1). Furthermore, defects were also found in the palates of Class 1 and 2 mutants. A small subset of these embryos had cleft palate (Class 1 (n=4/10) and Class 2 (n=2/2)) while most were missing the palatal process of palatine (Figure 1). The mandible or lower jaw, which uses Meckel’s cartilage as a template for its development was present in all Class 1 and 2 *Sf3b4* NCC mutants. However, the mandible of Class 2 mutants were less ossified than those of controls (Figure S3A). Furthermore, in Class 3 mutants, the mandibles found were dysmorphic and showed no obvious distal and proximal patterning (Figure 1).

**Figure 1:**
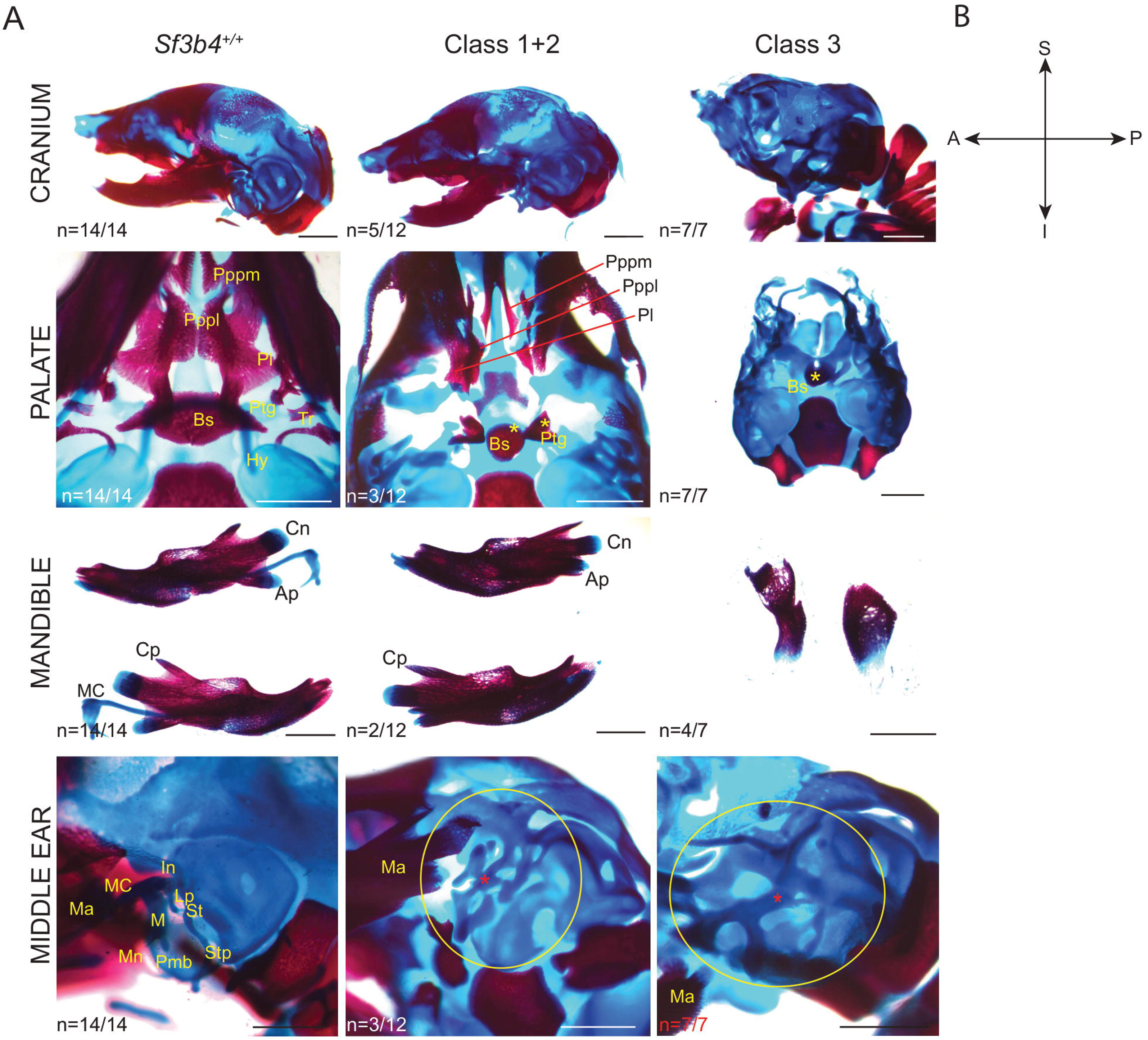
*Sf3b4* mutants at E17.5 have various skeletal craniofacial abnormalities. A. Craniofacial defects were seen in the cranium, palate, mandible and middle ear structures in *Sf3b4* NCC mutants. An: angular process, Bs: basisphenoids, Cn: condylar process, Cp: coronoid process, Hy: Hyoid, In: Incus, Lp: lenticular process, M: malleus, Ma: mandible, MC: Meckel’s cartilage, Pbm: processus brevis malleus, Pl: palatine, Pppl: palatal process of palatine, Pppm: palatal process of premaxilla, Ptg: Pterygoid, St: stapes, Stp: styloid process. red * indicates dysmorphic middle ear cartilage. Sale bar is 1000μm. B. Schematic to indicate orientation of the cranium. A: anterior, I: inferior, P: posterior, S: superior.

In wildtype embryos, the proximal/distal portion of the Meckel’s cartilage contributes to the malleus and incus - two of three ossicles of the middle ear. The third middle ear ossicle, the stapes, is derived from NCCs and mesoderm, and forms independently of Meckel’s cartilage. In *Sf3b4* NCC mutants the proximal portion of Meckel’s cartilage was bilaterally reduced in Class 1 mutants (n=2/10) (Figure 1) and missing in all Class 3 mutants. Similarly, middle ear ossicles formed from this cartilage, were malformed or missing. The most severe defects were seen in Class 3 mutants, which lacked all middle ear ossicles and had ectopic cartilages in their places (Figure 1). Also, a subset of Class 1 and Class 2 mutants lacked the lateral process of the malleus and had hypoplasia of the manubrium and the lenticular process of the incus (n=3/12, Figure S3B). Additionally, the stapes was absent in a subset of Class 1 and Class 2 mutants. Interestingly, the middle ear ossicles are derived from the 1^st^ and 2^nd^ PA which were hypoplastic at earlier embryonic stages.

*Sf3b4* NCC mutants also had abnormalities in neural crest derived cartilages and bones of the neck. The clavicles were asymmetric, shorter and dysmorphic in Class 1 (n=3/10), Class 2 (n=1/2) and all Class 3 (n=7/7) mutants (Figure S3C). In one Class 1 mutant, the body of the hyoid was not ossified, it was vertically positioned and abnormally attached to the cricoid cartilage (Figure S3D). In another, the hyoid was malformed and attached to the thyroid cartilage as two separate horn-like structure (Figure S3E). The hyoid was missing in all severely affected mutants. Additionally, the thyroid cartilage was dysmorphic in Class 1 embryos (n=1/10) (Figure S3E), and along with the tracheal rings were also missing in Class 1 (n=1/10) (Figure S3D), Class 2 (n=1/2) and all Class 3 (n=7/7) mutants. These analyses indicate that *Sf3b4* NCC embryos, including normal appearing Class 1 mutants have mineralization, cartilage, and bone defects. Our data also shows that mouse *Sf3b4* is required for proper morphogenesis of NCC-derived cartilages and bones that are malformed in patients with SF3B4-related syndromes.

### *Sf3b4^ncc/-^* embryos have vertebral transformations and malformations

We previously reported vertebral transformations and missing vertebrae in E17.5 *Sf3b4^+/–^* embryos^21^, thus, we analyzed the axial skeleton of E17.5 *Sf3b4* NCC mutants for defects. Most *Sf3b4^ncc/ncc^* mutant embryos belonging to Class 1 (n=5), Class 2 (n=2) and Class 3 (n=1) were indistinguishable from their control littermates. However, one *Sf3b4^ncc/ncc^* Class 1 embryo had an ectopic rib at C7 that joined to the first rib at T1, while another embryo of the same genotype belonging to Class 3 had the 8^th^ rib bilaterally connected to the sternum. On the other hand, axial abnormalities like those found in *Sf3b4*^+/–^ embryos were observed in all *Sf3b4^ncc/–^* mutant embryos. These abnormalities included the presence of ectopic rib at C7 attached to the rib at T1 (n=6), in 46% of embryos(Figure S3F), T1 to T2 transformation in 7% of embryos (n=1), and a missing L6 in all mutants (n=12). The type and incidence of vertebral malformations found in *Sf3b4^ncc/–^* embryos are similar to what was previously reported, consistent with the requirement of normal levels of Sf3b4 in the mesoderm.

### *Sf3b4* is required for cranial ganglia formation

NCC along with ectodermal placodes give rise to the cranial ganglia and their sensory neurons^29^. Immunohistochemistry using the neurofilament 2H3 antibody revealed missing or abnormally formed cranial sensory ganglia in E10.5 *Sf3b4* NCC mutant embryos. In Class 1 mutant (n=1/7), cranial nerve (CN) V, VII and X were indistinguishable from those of controls. Whereas in Class 2 (n=1/7) and Class 3 (n=5/7) mutants, the distal ends of CN V – ophthalmic, maxillary, and mandibular branches – (Figure 2B, arrow) did not extend as far ventrally as in controls. In addition, the proximal and distal regions of CN V were not connected in Class 3 mutants (Figure 2C, arrow). More caudally, whereas CN VII and CN IX failed to extend as far ventrally as in controls in the Class 2 mutant (Figure 2B), CN IX and CN XII were not detected in Class I and Class 2 mutants, respectively (Figure 2B, white star). Furthermore, in Class 3 mutants, CN IX and CN XII did not extend as far proximally and distally as they did in controls (Figure 2C). Additionally, while the dorsal root ganglia were reduced in the rostral region and completely absent in the caudal end of the Class 2 mutant, in three Class 3 mutants, all sensory ganglia were severely reduced. Altogether, these findings indicate that loss of *Sf3b4* in NCC disrupts development of NCC-derived ganglia.

**Figure 2:**
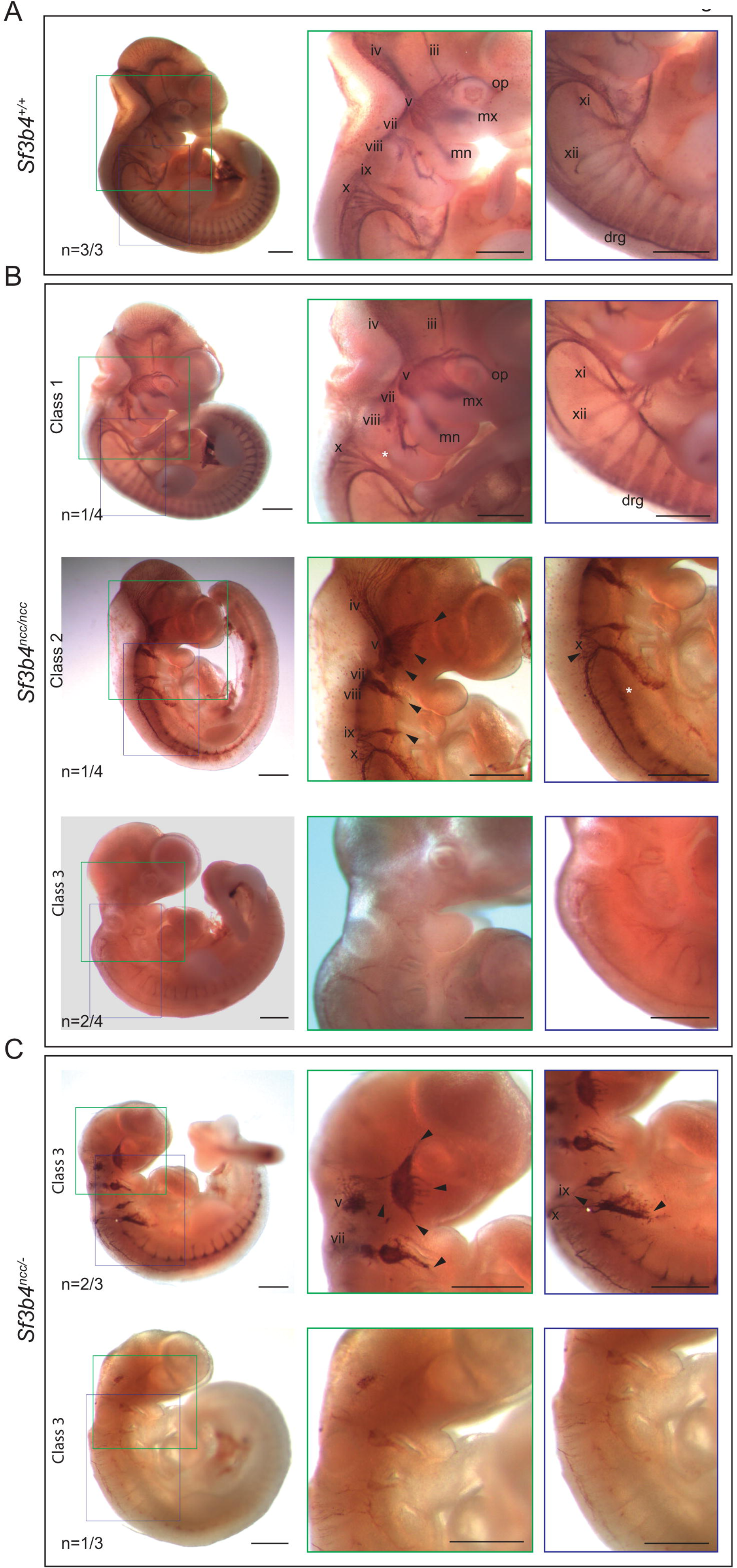
Normal development of cranial sensory ganglia in E10.5 embryos is affected by mutations in *Sf3b4*. Cranial neurons are numbered, mn: mandibular, mx: maxillary, op: ophthalmic, drg: dorsal root ganglion. High magnification is shown beside the embryos with the colour of the box corresponding to the area of the high magnification image. Black arrows indicate the incomplete formation of the cranial nerves. White star indicated missing cranial nerves. Scale bar is 500μm. A. Representative image of 2H3 staining in wild type embryos B. Images of 2H3 staining in *Sf3b4^ncc/ncc^* embryos from Class 1 (top panel), Class 2 (middle panel) and Class 3 (bottom panel). In these embryos, cranial nerves are either absent or incomplete. C. Images of 2H3 staining in *Sf3b4^ncc/-^* embryos (all Class 3) showing reduced and discontinuous cranial nerves or complete absence of neurofilament.

### *Sf3b4* is required for normal heart development

In addition to the cartilage and skeletal structures of the head and face, NCCs also contribute to the developing heart. Cardiac NCCs migrate into the outflow tract of the developing heart and contribute to the aorticopulmonary septum and the endocardial cushion^30^. By E12.5, septation of the distal outflow tract has completed^31^. Additionally, we observed a large increase in the percentage of dead/reabsorbed *Sf3b4* NCC mutants at this stage (Sup Table 1.1 and 1.2). To determine if heart malformations contribute to the significant loss of *Sf3b4* mutants at birth, E12.5 embryos were analyzed. No discernable differences were found in wholemount hearts of controls (n=2) and Class 1 *Sf3b4^ncc/ncc^* (n=2) mutant embryos (Figure 3A and 3B). In contrast, a similar analysis of the hearts of Class 2 *Sf3b4^ncc/ncc^* (n=2) and Class 3 *Sf3b4^ncc/–^* (n=2) mutants showed that the right and left atria were hypoplastic and that the aorticopulmonary septum, which separates the aortic and the pulmonary trunks, was not present (Figure 3C). However, histological analysis of frontal and sagittal sections of E12.5 embryos revealed heart defects in *Sf3b4* NCC embryos of all Classes. In Class 1 mutant embryos (n=2), the aorticopulmonary septum (Figure 3E’ and 3F’) was thicker than those of controls, whereas walls of both ventricles were thicker in mutants of all classes (n=6) (Figure 3F’, G’ and H’, black line). Additionally, the dorsal aorta was smaller than those of controls in Class 1 mutants (Figure 3E’’ and 3F’’), abnormally connected to the pulmonary trunk in Class 2 mutants (Figure 3G’’), and undetectable in Class 3 mutants (Figure 3H”). Similarly, the aorticopulmonary trunk was not found in Class 2 and Class 3 mutants (Figure 3G’ and H’) leading to the presence of a common truncus. Additional heart defects seen in Class 3 *Sf3b4^ncc/–^* mutants included a smaller heart, that did not completely occupy the mediastinum, a small and abnormally positioned left and right atriums, and an absence of the aortic/pulmonary valves (Figure 3H’), when compared to wild type (Figure 3E’). Thus, *Sf3b4* NCC mutants have several heart defects that have been reported in patients with SF3B4-related syndromes^5^ and reveal a role for SF3B4 during septation of the outflow tracts as well as in the differentiation and growth of the atria and ventricles. Based on these findings, we infer that cardiac abnormalities may contribute to loss of *Sf3b4* NCC mutants.

**Figure 3:**
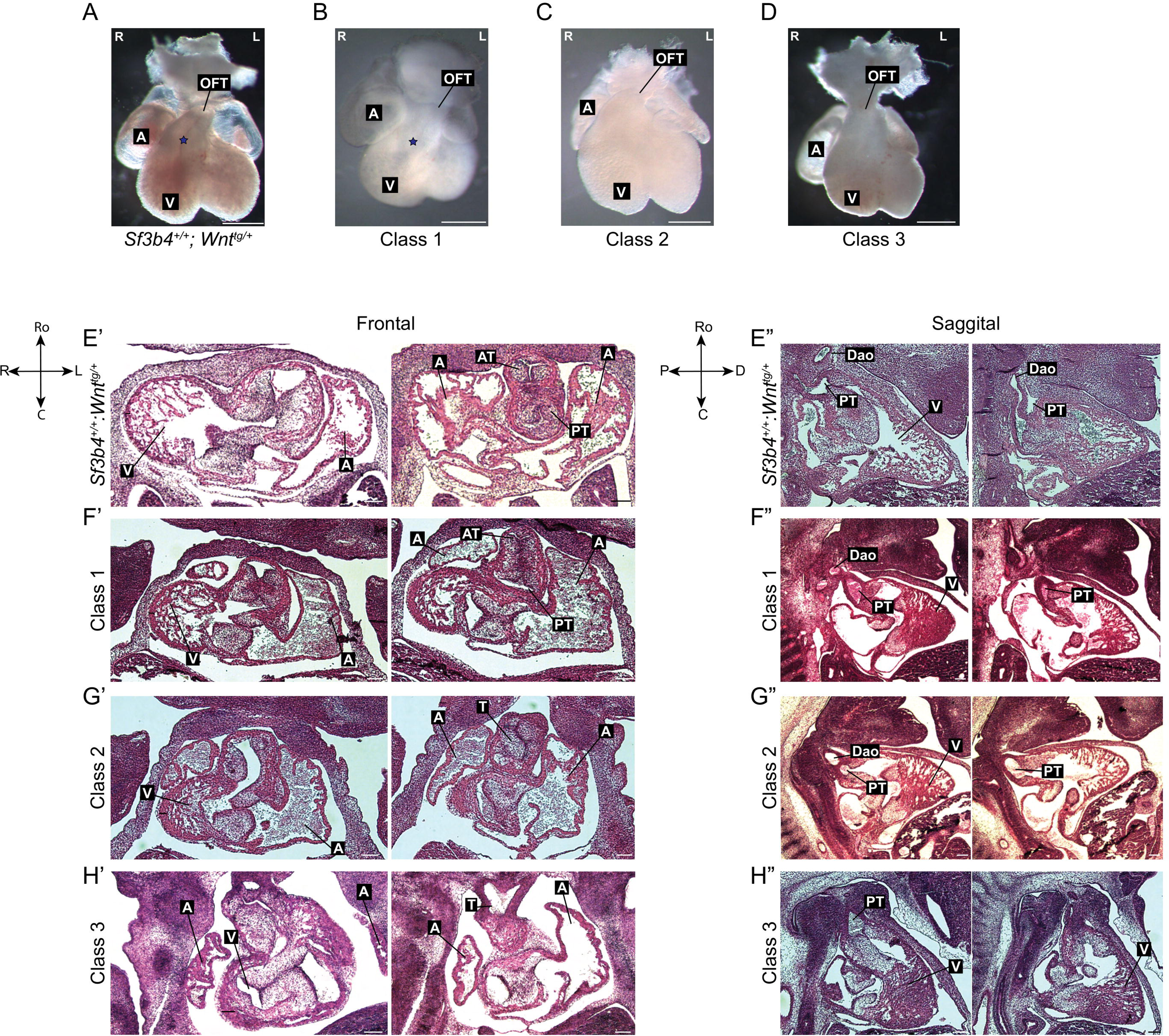
Heart defects were observed in E12.5 *Sf3b4* mutants. A-D. Whole mount heart of wild type and mutants. A. *Sf3b4^+/+^;Wnt^tg/+^*, B. Class 1 *Sf3b4^ncc/ncc^*, C. Class 2 *Sf3b4^ncc/ncc^* with normal craniofacial region and hypoplastic mid- and hindbrain, D. Class 3 *Sf3b4^ncc/-^*. A: atrium, L: left, OFT: outflow tract, blue star indicates the visible aorticopulmonary septation in the outflow tract. Scale bar is 1000um. For all frontal sections, orientation is indicated beside E’. C: caudal, L: left, R: Right, Ro: rostral. Black line shows thickness of the ventricle wall. For all sagittal sections, orientation is indicated beside E”. C: caudal, D: distal, Dao: dorsal aorta, P: proximal, Ro: rostral. A: atrium, AT: aortic trunk, PT: pulmonary trunk, V: ventricle. Scale bar is 100μm. E’. Frontal, serial sections of heart in a wild type embryo, showing formation of the aortic and pulmonary trunks. E”. Sagittal, serial sections of heart in a wild-type embryo showing the dorsal aorta arising from the left ventricle. F’. Frontal, serial sections of heart in a Class 1 *Sf3b4^ncc/ncc^* embryo, showing thickened aortic and pulmonary trunks, smaller right atrium and thicker ventricle wall. F”. Sagittal, serial sections of heart in a Class 1 *Sf3b4^ncc/ncc^* embryo, showing formation of a small dorsal aorta. G’. Frontal, serial sections of heart in a Class 2 *Sf3b4^ncc/ncc^* embryo, showing formation of a single truncus and thereby absence of aorticopulmonary septum, small right and left atrium and thicker ventricle wall G”. Sagittal, serial sections of heart in a Class 2 *Sf3b4^ncc/ncc^*, showing dorsal aorta fusing with the pulmonary trunk in the right ventricle instead of arising from the left ventricle. H’. Frontal, serial sections of heart in a Class 3 *Sf3b4^ncc/-^* embryo showing a small heart with small right and left atrium, thicker ventricle wall and single truncus with no valves. H”. Sagittal, serial sections of heart in a Class 3 *Sf3b4^ncc/-^* embryo showing absence of the dorsal aorta.

### Fewer *Sf3b4* mutant NCC are found in craniofacial region due to reduced proliferation and increased cell death

Hypoplasia and absence of cranial neural crest cell derivatives in *Sf3b4* mutants could be due to reduced NCC generation, migration or survival. To track CRE expressing NCC, we used the *Rosa^mT/mG^* reporter, which marks these cells and their derivatives with GFP^32^. At E8.5, the distribution of GFP+ NCC found in the head and in streams at the levels of the 1^st^ and 2^nd^ pharyngeal arches of *Sf3b4^ncc/ncc^* mutants and controls were comparable (Figure 4A). However, the GFP signal appeared patchy and reduced in the frontonasal region of mutants (Figure 4A, arrow). Furthermore, few GFP+ cells were found in the 1^st^ pharyngal arch and frontonasal prominences of a phenotypically normal E8.5 *Sf3b4^ncc/–^* mutant with 8-somites (Figure 4A), when compared to a somite-matched control (Figure 4A and Figure S4A). A day later, significantly less GFP+ cells were found in the 1^st^ pharyngeal arch of E9.5 *Sf3b4* NCC mutant embryos (t-test; *Sf3b4^ncc/ncc^* p=0.0006, *Sf3b4^ncc/–^* p=0.0001) (Figure 4B, and 4C). We postulate that the reduced number of GFP+ (*Sf3b4* homozygous mutant) cells found in the 1^st^ pharyngeal arch and frontonasal prominence of E8.5 *Sf3b4* NCC embryos is due to a reduction in *Sf3b4* NCC generation, as previously shown in *Xenopus* Sf3b4 morphants^24^, as well as reduced cell survival in the developing craniofacial region.

**Figure 4:**
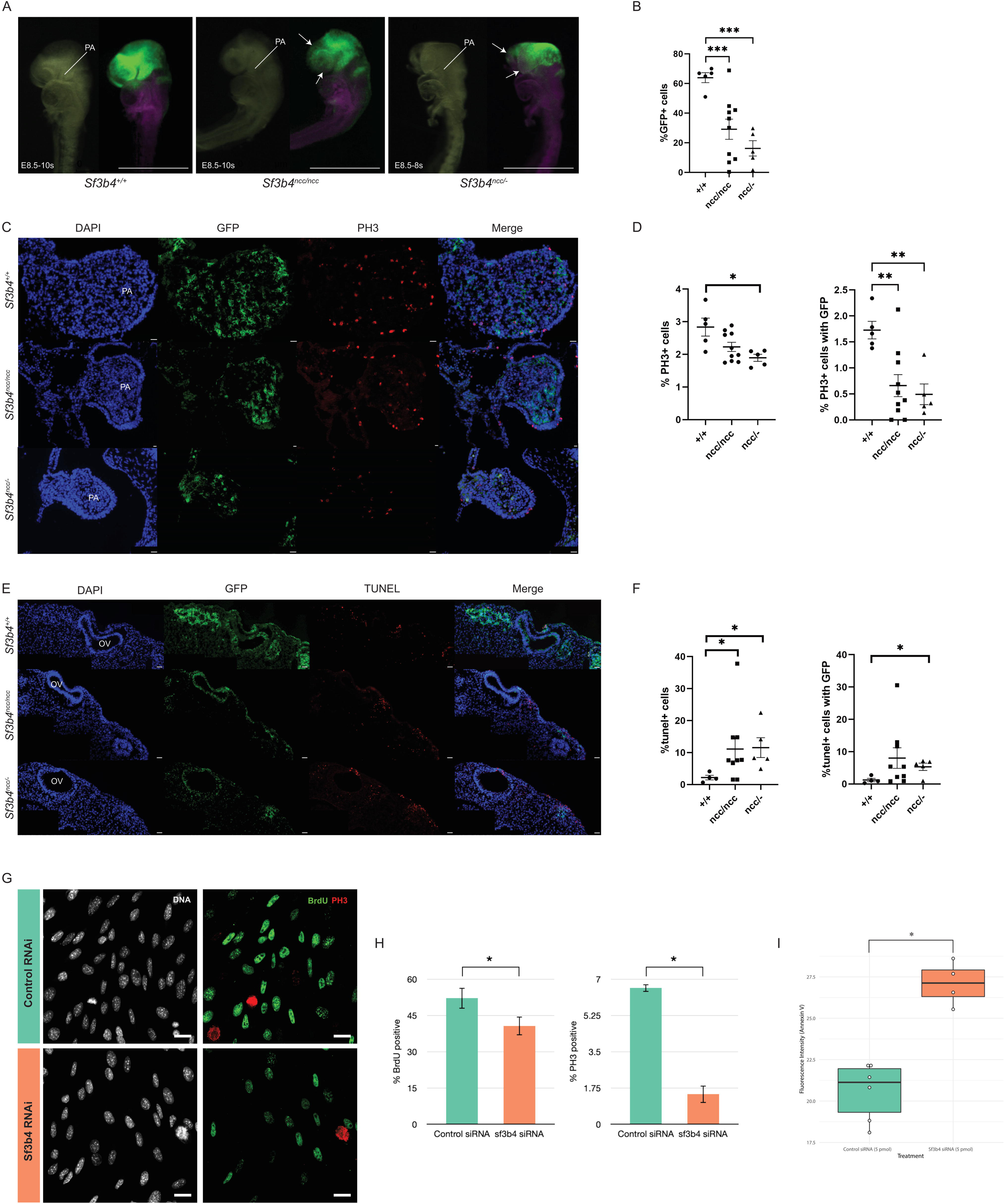
NCCs with *Sf3b4* mutations have decreased proliferation and increased cell death. A. *Sf3b4^+/+^;Wnt^tg/+^* embryos with R^mT/mG^ have GFP expression in the head and 1^st^ pharyngeal arch where as decreased GFP expression is observed in the frontonasal region of *Sf3b4^ncc/ncc^* embryos, white arrow. And *Sf3b4^ncc/-^* embryos have decreased GFP in the head, white arrow, and no GFP in the 1^st^ PA. Images are shown in brightfield followed by fluorescence. PA; pharyngeal arch Scale bar is 20μm. B. Quantification of GFP expression in the 1^st^ PA at E9.5 with significant decrease in *Sf3b4* mutants. Each data point represents the average of 4 sections from one embryo, error bars are ±SEM, ***p<0.001. C. Representative images of 1^st^ pharyngeal arch sections of wild type and mutant embryos stained with PH3 to look at proliferation. PA; pharyngeal arch, Scale bar is 20μm. D. Graph showing quantification of PH3+ cells in the pharyngeal arch with significant decrease in the mutants. Each data point represents the average of 4 sections from one embryo, error bars are ±SEM, *p<0.05, **p<0.005. E. Representative images of TUNEL assay on sections of hindbrain with otic vesicle in wild type and mutant embryos. OV; otic vesicle. Scale bar is 20μm. F. Graph showing quantification of TUNEL+ cells in the hindbrain and otic vesicle region with significant increase in the mutants. Each data point represents the average of 3/4 sections from one embryo, error bars are ±SEM, *p<0.05, **p<0.005. G. BrdU and PH3 staining of O9-1 cells treated with control or Sf3b4 siRNA. Scale bar is 10μm. H. Quantification of BrdU+ and PH3+ cells showing significant decrease in cells treated with *Sf3b4* siRNA. *p≤0.05. I. Quantification of Annexin V fluorescence intensity in control (n=6) and knock down (n=4) samples. A Kruskal-Wallis rank sum test was used to determine the significance associated with the difference between the treatment groups. *p≤0.05.

Cell proliferation and cell death was measured in the E9.5 embryos to determine if reduced proliferation and/or increased cell death contribute to the reduced number of GFP+ cells found in the 1^st^ pharyngeal arch of *Sf3b4* NCC mutants. Histone H3 phosphorylation on serine-10 staining was used to label mitotic cells in E9.5 control (n=5), and *Sf3b4* NCC embryos [*Sf3b4^ncc/ncc^* (n=10, 5 each from Class 1 and Class 3) and *Sf3b4^ncc/–^* (n=5, all Class 3) embryos]. In the 1^st^ pharyngeal arch, the proportion of GFP+ cells stained with PH3 was significantly reduced in *Sf3b4^ncc/ncc^* and *Sf3b4^ncc/–^* mutants, when compared to controls (t-test, p=0.0018 and p=0.0016, respectively) (Figure 4C and 4D). However, the proportion of all cells (GFP+ and GFP-) positive for PH3 was only significantly reduced in 1^st^ pharyngeal arches of *Sf3b4^ncc/–^* mutants, when compared to controls (t-test, p=0.0233) (Figure 4C and 4D). To quantify the number of cells undergoing apoptosis, the proportion of cleaved caspase-3 positive cells found in the in the 1^st^ pharyngeal arch of controls (n=5), and *Sf3b4* NCC mutants (*Sf3b4^ncc/ncc^* (n=10) and *Sf3b4^ncc/–^* (n=5)) was counted. However, this analysis revealed no difference between *Sf3b4* NCC mutants and controls (Figure S4F). To more broadly label dying cells, TUNEL was used. No difference was found in the proportion of TUNEL-positive cells in the 1^st^ pharyngeal arch of controls (n=4), *Sf3b4^ncc/ncc^* (n=9), and *Sf3b4^ncc/–^* mutants (n=5, Figure S4D and S4E). This data shows that reduced proliferation contributes to the reduced number of NCCs found in the 1^st^ pharyngeal arch of *Sf3b4* NCC mutants. Our findings also suggest that reduced proliferation of GFP^−^ *Sf3b4* heterozygous cells in the 1^st^ pharyngeal arches of *Sf3b4^ncc/–^* may underlie the more severe phenotypes found in *Sf3b4^ncc/–^* embryos.

We performed a similar analysis in the region immediately surrounding the hindbrain and otic vesicle, where the NCC emerge, to determine if *Sf3b4* mutant NCCs outside of the 1^st^ pharyngeal arch shows similar defects in survival. This analysis revealed no difference in the proportion of PH3-positive found in wild type and *Sf3b4* NCC mutants (Figure S4B and S4C). In contrast, the proportion of GFP+ cells that were also TUNEL positive was significantly increased in *Sf3b4^ncc/–^* mutants (t-test, p=0.018) (Figure 4E and 4F). Additionally, the total number of cells (GFP+ and GFP–) positive for TUNEL was significantly increased in *Sf3b4* NCC mutants when compared to controls (t-test, *Sf3b4^ncc/ncc^* p=0.043 and *Sf3b4^ncc/–^* p=0.037) (Figure 4E and 4F). These data indicate that increased cell death contributes to reduced NCC survival outside of the 1^st^ pharyngeal arch and shows that *Sf3b4* homozygous mutant NCCs have a greater propensity to die if the neighboring cells are heterozygous for this gene.

To further investigate the impact of reduced *Sf3b4* specifically to NCC proliferation and survival, we investigated Sf3b4 knock-down in O9-1 cells, a mouse neural crest cell line. We found that the proportion of BrdU and PH3 positive O9-1 cells was significantly reduced in cells transfected with *Sf3b4* siRNA compared to control transfections (Figure 4G and 4H). Additionally, to investigate NCC death after *Sf3b4* knockdown, O9-1 cells were stained with a fluorophore-conjugated Annexin V 24 hours post-transfection. Annexin V binds to membrane phosphatidylserines that are exposed to the extra-cellular environment during the early stages of apoptosis^33^. The Annexin V assay revealed that O9-1 cells transfected with *Sf3b4* siRNAs fluoresce at significantly higher levels relative to the control condition (Figure 4I). These data indicate that normal levels of *Sf3b4* are necessary for NCC proliferation and survival both *in vivo* and *in vitro*.

### Homozygous deletion of *Sf3b4* disrupts the transcriptomic landscape before the onset of morphological defects

*Sf3b4* is important for the U2 snRNP to attach upstream of the branch point site^17^, we next compared the transcriptome of E8.5 normal looking *Sf3b4^ncc/–^* and wild type embryos between 8-10s using RNAseq. Previously, a similar analysis of E9.5 *Sf3b4^+/–^* embryos and somites showed an increase in exon skipping associated differential splicing events (DSE) and few differentially expressed genes^21^. In contrast, RNAseq analysis of E8.5 *Sf3b4* NCC mutants revealed 733 differentially expressed genes, of which 330 genes were significantly upregulated, and 403 genes were significantly down-regulated (Figure 5A). Additionally, a greater than 5-fold increase in skipped exon (SE) events was found in *Sf3b4* NCC mutants, when compared to controls; 315 SEs were found in 246 genes in control, versus 1,620 SEs in 1,020 genes in mutants (Figure 5B and Figure S5A). To determine if DSEs found in *Sf3b4^ncc/-^* mutants were associated with any molecular function or biological processes, gene ontology and KEGG pathway were analysed. No enrichment was found for any splice events except for one term, H4K20 histone methyltransferase activity, for alternative 5’ splice site (A5SS) events (Figure S5B). Thus, homozygous deletion of *Sf3b4* in NCC alters the transcriptomic landscape of mutant embryos prior to any morphological defects.

**Figure 5:**
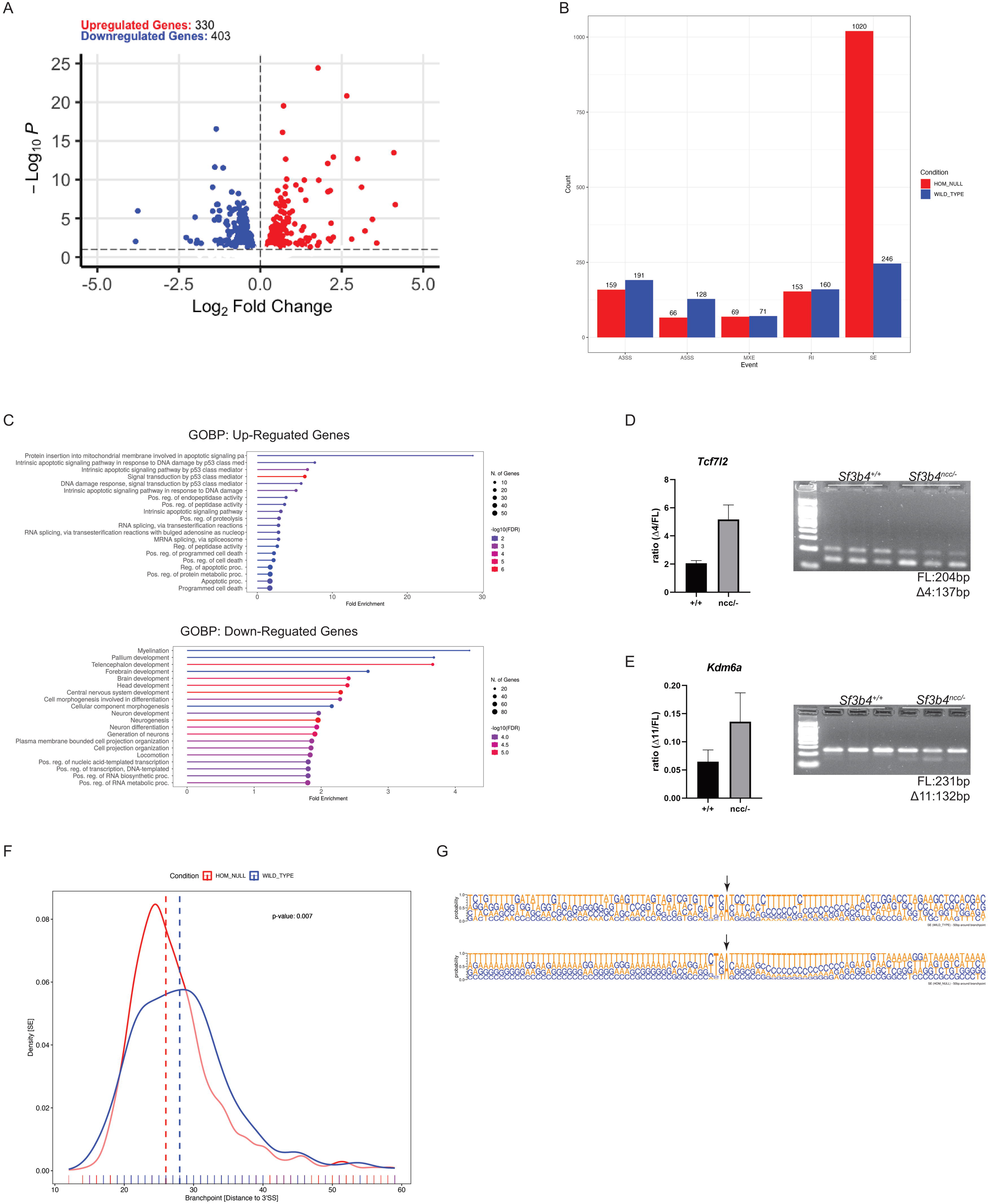
RNAseq reveals several differentially expressed and differentially spliced genes. A. Volcano plot showing DEG with padj<0.05. B. Graph showing number of genes that were differentially spliced for each of the splicing events. C. Gene ontology biological processes of DEGs that are upregulated (top panel) and down regulated (bottom panel). D. RT-PCR showing increased skipping of exon 4 in *Tcf7l2* (left panel) and quantification of gel (right panel). E. RT-PCR showing increased skipping of exon 11 in *Kdm6a* (left panel) and quantification of gel (right panel). F. Predicted branch point signals for SE events with dashed line showing preference for branch point at specific distance from 3’SS in wildtype (28bp) and mutant (26bp). G. Predicted sequence composition 50bp around the branch point showing an increase in thymine bases surrounding the branch-point A in mutants.

### The expression levels of genes in the NCC regulatory network are reduced in *Sf3b4* NCC mutants

Transcripts in the NCC gene regulatory network^34^ important for NCC development were mostly decreased in *Sf3b4* NCC mutants. Specifically, we found significant reduction in expression of *Tfap2b* part of the neural plate border module, and in several *Sox* genes important in NCC specification and migration: *Sox5*, *Sox6*, *Sox9* and *Sox10* (Sup Table 5). Several other genes important for neural crest generation were also significantly reduced in mutants, including: *Ets1*, *En2* and *Grhl3*, *Dnmt3a*, *Dnmt1*, and *Nes* (Sup Table 5). In contrast, *E-cadherin* levels were significantly increased in these mutants. E-cadherin is downregulated when cells undergo epithelial-mesenchymal transition (EMT), an essential first step in neural crest migration from the hindbrain. Furthermore, HCR revealed reduced levels of *Tfap2b* and *Sox9* along with uneven patterning of *Sox9* in the neural plate of morphologically normal E8.5 *Sf3b4^ncc/ncc^* mutants (n=3) with 7/8s when compared to somite matched controls (Figure S5C).

To identify pathways dysregulated in *Sf3b4* NCC mutants, gene ontology analysis using all significantly differentially expressed genes was performed. Several pathways were enriched in these mutants, including intrinsic apoptotic signalling pathway by P53 amongst upregulated genes, and neurogenesis amongst the most downregulated (Figure 5C). Based on these data, we hypothesized that NCC with reduced *Sf3b4* levels may arrest during the early stages of EMT and undergo P53-mediated cell death.

### Reducing levels of *Trp53* did not rescue malformations or viability of *Sf3b4* NCC mutant embryos

Increased P53 activity contributes to craniofacial malformations in several mouse models of neurocristopathies and spliceosomopathies^27,28,35,36^, while perturbation of global molecular processes that activate P53 contributes to neural crest cell death in human cell lines and in mouse models^35^. Therefore, we next sought to determine if P53-activity was increased in cells with reduced levels of *Sf3b4.* RNAseq analysis showed significant enrichment of the P53 pathway, and significant increases in the levels of P53-downstream targets: *Mdm2* (1.73-fold), *Phlda3* (21.75-fold), *Trp53inp1* (7.41), *Ccng1* (6.35-fold), *Cdkn1a* (20.96-fold) and *Eda2r* (17.04-fold). In addition, when P53 activity was analyzed in O9-1 cells transfected with *Sf3b4* siRNA, we found a remarkable increase in activated P53 (Figure S6A). To evaluate the extent of P53 activation upon *Sf3b4* knockdown in O9-1 cells, the expression of cell cycle genes that are directly and indirectly regulated by P53 was quantified by qRT-PCR. This analysis revealed a significant upregulation of *Ccng1* (2.63-fold) and *Cdkn1a* (3.23-fold) in knockdown samples relative to control (Figure S6B). These data reveal that *Ccng1* and *Cdkn1a*, two important mediators of the G1-S phase transition^37–39^ that are transcriptionally regulated by P53 are significantly overexpressed in knockdown samples compared to control. We further evaluated whether *Sf3b4* knockdown leads to activation of the P53 apoptosis pathway. qRT-PCR data showed a significant upregulation of P53-regulated genes, including *Mdm2* (5.61-fold), *Phlda3* (2.78-fold), *Puma* (3.72-fold), and *Trp53inp1* (11.73-fold) in *Sf3b4* knockdown cells compared to control (Figure S6C), suggesting a role for P53 mediated apoptosis in NCC with loss of *Sf3b4*.

Sakai and colleagues showed that neuroepithelial cells in E8.5 mouse embryos have higher levels of ROS relative to other cells and that exposure of wild-type mouse embryos to a ROS generator drug (3-NP) leads to a substantial increase in neuroepithelial cell death^40^. The marked upregulation of *Trp53inp1*, a gene with known antioxidant function, in *Sf3b4* knockdown cells (Figure S6C) led to the consideration of the cells’ oxidative state as a factor that might influence the observed increase in apoptosis. To evaluate this hypothesis, the levels of ROS and Superoxide were assayed in *Sf3b4* knockdown and control O9-1 cells. Interestingly, while the levels of both ROS and Superoxide for the control wells clustered together, the levels for the knockdown wells were much more variable (Figure S6D and S6E). To further evaluate the role of oxidative stress in *Sf3b4*-mediated apoptosis, the expression of key antioxidant genes *Sod1*, *catalase*, and *Nqo1*^41^ was also assayed in *Sf3b4* knockdown and control cells (Figure S6F and 6G). Although the group means for each gene were not significantly different, their reference-normalized expression (ΔCT) for the control condition exhibited higher variance relative to the knockdown condition. Taken together, and assuming a consistent transfection efficiency across all wells, these data suggest that *Sf3b4* knockdown creates an overall more stressful environment for NCCs.

Reducing levels of P53 has variable impact on craniofacial development and survivability of mouse models of craniofacial spliceosomopathies. For *Snrpb* NCC mutants, reducing levels of P53 resulted in increased survival but failed to rescue patterning defects in the mandible^27^. In contrast, reducing levels of P53 alone, proved insufficient to improve craniofacial development or survival of *Eftud2* NCC mutants^27,42^. To test if decreasing P53 rescues the craniofacial malformations seen in *Sf3b4^ncc/–^* mutants, we mated *Sf3b4^+/–^*;*Trp53^L/+^;Wnt1-cre2^tg/+^* mice with *Sf3b4^L/+^*;*Trp53^L/L^* mice to generate pups that have homozygous deletions for both *Sf3b4* and *P53*. We followed pups from 4 litters from P0 to P21. Of 42 pups born from these matings, four died at P1 and could not be followed to weaning. However, genotyping of carcasses collected from these pups revealed that two were double heterozygous mutant for *Sf3b4* and *Trp53* (*Sf3b4^L/+^;Trp53^L/+^;Wnt1-Cre2^tg/+^*) and 2 were conditional mutants that did not carry CRE (*Sf3b4^L/+^;Trp53^L/+^*). Furthermore, zero *Sf3b4* and *Trp53* double homozygous NCC mutants was found amongst the 38 pups weaned at P21. Additionally, E9.5 *Sf3b4* and *Trp53* double homozygous NCC mutants were indistinguishable from Class 3 E9.5 *Sf3b4* NCC mutants (Figure S6H). Thus, we conclude that increased P53 activity is not sufficient to explain the observed malformations and reduced survival associated with loss of *Sf3b4* in NCC.

### Mis-splicing of chromatin remodelers contributes to malformations in *Sf3b4* mutants

Previously, histone modifiers were found to be differentially spliced in *Sf3b4^+/–^* embryos when compared to controls^21^. Consequently, a list of genes with significant increases in SE with FDR<0.05 was used to query MGI for gene ontology terms related to histone modifier and chromatin remodeler, and craniofacial and heart development. Seven genes: *Tcf7l2*, *Kdm6a*, *Kdm5c*, *Setd5*, *Carm1*, *Dnmt3a* and *Dnmt2b* were identified from this analysis (Table1). Next, RT-PCR was used to show that SE events predicted in *Tcf7l2* and *Kdm6a* was also seen in RNA collected from new pools of *Sf3b4* mutants (Figure 5D and 5E). In addition, RT-qPCR revealed an increase in levels of *Kdm5c*, *Carm1* and *Setd5* in E8.5 *Sf3b4* mutant samples, when compared to controls although not significant (Figure S5D). Thus, we concluded that reducing *Sf3b4* levels in NCC affects splicing of transcription factors and histone modifiers involved in normal craniofacial development.

### *Sf3b4* is necessary for proper splicing and branchpoint recognition

To investigate whether aberrant splicing events were characterized by specific sequences, we compared alternative events preferentially found in mutant samples, wild-type samples and a set of 1000 randomly chosen alternative splicing events. No difference was found for 5’ splice site (SS) and 3’SS strengths of any alternative splicing events (skipped exons (SE), retained introns (RI), mutually exclusive exons (MXE), alternative 3’ splice sites (A3SS) and alternative 5’ splice sites (A5SS), although a trend towards weaker 5’SS strength was seen for SE, RI and A3SS events found in mutants (Figure S7B, S7C and S7D). We also looked at general base composition in all AS events. Although there was no significant difference found, the GC content was slightly decreased in SE and MXE events found in mutants (Figure S7E and S7F). We also analyzed the strength and position of predicted branch point (BP) signals and found a significant preference for a more proximal BP location in mutant specific SE events (26bp in mutants compared to 28bp in controls, p=0.007) (Figure 5F and Figure S7A). Furthermore, the sequence composition 50bp around the BP showed an increase in thymine bases surrounding the branch-point A, when compared to controls (Figure 5G). We previously showed significantly stronger 5’SS and non-significant preference for a more proximal branch point in mutant-specific A3SS events^21^, suggesting that reducing levels of *Sf3b4* perturbs branch-point selection. Additionally, the current study suggests that an increase in thymidine bases around the branch point may contribute to the selection of a more proximal BP usage and differential splicing in *Sf3b4* NCC mutants.

## DISCUSSION

Craniofacial spliceosomopathies, like SF3B4-related syndromes and MFDM which is caused by pathogenic variants in *EFTUD2*, are also termed neurocristopathies since craniofacial defects are predominantly found in structures derived from NCCs. In this study, we showed that homozygous deletion of *Sf3b4* in NCCs results in craniofacial defects with variable expressivity mimicking those seen in patients. Loss of *Sf3b4* in NCCs results in decreased NCC specifier genes as well as reduced survival of mutant NCCs and surrounding cells. Furthermore, though the P53 pathway was significantly upregulated by loss of *Sf3b4* in NCCs and resulted in increased expression of P53 pathway genes, craniofacial defects and survival of *Sf3b4* mutants was not solely dependent on P53, similar to what was found for *Eftud2* NCC mutants^42^. Additionally, we found aberrant splicing of key histone modifiers, as well as altered branch point usage that may contribute to the global disruption of the splicing landscape found in mutants.

The penetrance and severity of the *Sf3b4* mutation were modified by the presence of one or both wild type alleles of *Sf3b4* in neighbouring cells. We also show that loss of this gene perturbs all stages of NCC development: generation, migration, expansion, and differentiation, regardless of genotype. In *Sf3b4^ncc/–^* embryos, several NCC specifier genes, particularly *Tfap2b* that is required early in NCC specification^43^, and *Sox9* which is expressed in pre-migratory NCCs^44–46^, were reduced. In addition, *Sf3b4* mutants have significant decrease of *Sox10*, which is switched on when NCCs begin epithelial mesenchymal transition^47–49^, and increased expression of *E-cad*, which is normally downregulated in cells undergoing EMT^50,51^. Taken together, this suggests, that *Sf3b4* mutant NCC precursors experience a delay, or block, in EMT. Furthermore, *Sf3b4* mutant NCCs that do exit the neural tube, undergo increased cell death, and have a block in proliferation upon reaching the pharyngeal arches. Similar findings using O9-1cells suggest that global disruption of SF3B4-mediated splicing triggers death and arrest of mutants. Our findings are aligned with those of *Devoto* et al^24^ who also found reduced NCC migration when *Sf3b4* was knocked-down in Xenopus, and those of Yamada et al^22^ who showed reduced proliferation in the forebrain of *Sf3b4^+/–^* mouse embryos. However, unlike Xenopus embryos with *Sf3b4* knockdown, *Sf3b4* NCC mutant mouse embryos do not show an expanded neural tube. In fact, their neural ectoderm was thinner than those of controls. We propose that this is likely due to a block of proliferation and/or increased cell death of NCC precursor cells that fail to exit the neural tube. Finally, our data support a role for *Sf3b4* in differentiation, as proposed by Marques et al^13^, and also reveals a requirement for this gene in patterning of the lower jaw.

We uncovered several potential malformations that may contribute to the reduced numbers of *Sf3b4* NCC mutants born, including a previously unknown role for *Sf3b4* in heart development. We propose that *Sf3b4* NCC mutants of all Classes may die as a consequence of palatal and/or heart defects, similar to other mouse models of craniofacial spliceosomopathies^27,28^. Since a subset of Nager Syndrome patients shows heart defects, including coarctation of the dorsal aorta and complex heart defects, the role of SF3B4 in cardiac NCCs and heart mesoderm needs further investigation.

We previously reported that heterozygous loss of *Sf3b4* results in increased exon skipping and intron retention of histone modifiers required for development and patterning of the vertebrae^21^. To that end, we postulated that reduction of *Sf3b4* in NCCs would also result in aberrant splicing of histone modifiers required for craniofacial development. In fact, homozygous loss of *Sf3b4* in NCCs had a much greater impact on splicing, such that gene ontology enrichment did not show enrichment for any specific pathway, except for H4-K20 methyltransferase in A5SS events. H4K20 methylation is associated with chromatin remodelling, cell cycle regulation and DNA repair^52–54^. Thus, abnormal splicing of enzymes important for H4K20 methylation may contribute to dysregulation of these cellular processes in *Sf3b4* NCC mutants, or alternatively, dysregulation in these processes contribute to abnormal H4K20 methylation. Since the current study does not indicate the direction of this interaction, future studies will need to explore this finding.

Nonetheless, RNAseq data reveal increased skipped exons in a number of genes important for craniofacial and heart development, including: exon24 of *Kdm5c*, exon11 of *Kdm6a* and exon 4 of *Tcf7l2 (Tcf4)*. SE events found in *Kdm5c* and *Kdm6a* are predicted to disrupt their protein interaction. Though *Kdm5c* and *Kdm6a* are X-linked, and we do not see a sex-specific modification of expressivity or penetrance to *Sf3b4* loss in mice, combined reduction of these two histone demethylases may interact to contribute to malformations seen in male and female *Sf3b4* mutants. Additionally, TCF4 binds to β-catenin and is a key transcription factor important for mediating Wnt signalling^55,56^. The reported skipping of exon 4 is predicted to result in loss of the CTNNB1 domain of TCF4 which is important for binding to β-catenin. Thus, we expect that expression of WNT target genes essential for craniofacial and heart development would not be activated. Hence, although these SE events were not significant by RT-PCR, likely a result of the reduced sensitivity of this method when compared to RNAseq, we postulate that mis-splicing of these and other histone modifiers, as well as *Tcf7l2* contribute to craniofacial and heart malformations seen in *Sf3b4* mutants.

Like previous analyses of spliceosomal NCC mutants^27,28^, mis-splicing in *Sf3b4* mutant NCCs was associated with increased DEG and DSE. However, in *Sf3b4* NCC mutants, we also found sequence-specific alterations. Specifically, significant usage of a more proximal branch point, not seen in *Eftud2* and *Snrpb* NCC mutants, is predicted to result in increased use of A3’SS^57^. Additionally, the increased thymidine context of the branchpoint adenine in these SEs may alter yet to be identified splicing regulatory elements necessary for exon and intron definition. Nonetheless, we predict that variations in splice site characteristics found in these mutants combine to contribute to differential splicing in specific sets of transcripts in a tissue-specific manner. Future studies using single cell RNAseq and spatial transcriptomics would shed light on this hypothesis. Altogether, our studies are starting to reveal the specific features that are altered by mutation of core splicing factors, and a level of splicing regulation that has remained an enigma.

Overall, we show that craniofacial malformations similar to those found in patients with *SF3B4-*related syndromes were present in the majority of *Sf3b4* NCC mutants. In addition to craniofacial defects, *Sf3b4^ncc/–^* embryos also had vertebral abnormalities like those seen in *Sf3b4^+/–^* embryos^21,22^. Therefore, the *Sf3b4* NCC mutant embryos described herein are the first to model the spectrum of malformations and expressivity found from Nager to Rodriguez syndrome ^1,3^.

## Acknowledgements

We would like to thank Dr Mitra Cowan, Platform Manager, McGill Integrated Core of Animal Modeling for performing the microinjection experiments. We would like to thank the McGill Genome Centre for running the RNAseq. We also thank members of the Majewska laboratory their helpful comments on the manuscript. We thank the Research Institute of McGill University Health Centre for supporting S.K. We would also like to acknowledge the professional and technical support from the Animal Resource Division of RI-MUHC for maintaining our mice colonies.

## Contributions

S.K. carried out and analyzed all of the mouse and RNAseq experiments, drafted the manuscript and designed the figures. E.B. performed all initial RNAseq data analysis and generated part of the corresponding figures. E.C. performed part of the analysis on the RNAseq data and generated part of the corresponding figure. J.L. performed the IF experiments and part of the analysis. F.M., E.E.S., S.M. performed and analysed the O9-1 experiments. J.F. conceptualized and supervised O9-1 experiments and analysed and wrote the O9-1 sections of the manuscript. J.M. , L.A.J.-M. devised the project, L.A.J-M. conceptualized and supervised all the mouse experiments and wrote the manuscript.

## Funding

This project was funded by the Canadian Institutes of Health Research grant (202203PJT-480346-DEV-CFAC-157303) and by an Award by the Azrieli Foundation to Loydie Jerome-Majewska. The funders had no role in the study design, data collection and analyses, decision to publish or preparation of the manuscript.

## Declarations of interest

none

## KEY RESOURCES TABLE

| REAGENT or RESOURCE | SOURCE | IDENTIFIER |
| --- | --- | --- |
| **Antibodies** |  |  |
| PhosphoHistone H3 (PHH3) | Millipore | 06-570 |
| Cleaved Caspase-3 | Cell Signalling | 9661T |
| Alexa Fluor 568 | ThermoFisher | A-11001 |
| Neurofilament 2H3 | DSHB | N/A |
| HRP conjugated Anti-mouse IgG | Jackson ImmunoResearch | 115-035003 |
| **Chemicals, peptides, and recombinant proteins** |  |  |
| Alcian Blue | Therme Fisher | J60122.09 |
| Alizarin Red | Sigma | A5533 |
| DAB | Vector Laboratories | SK-4105 |
| Fluoroshield | Sigma | F6057 |
| RNA later | Invitrogen | AM7020 |
| iScript™ Reverse Transcription Supermix | Bio-Rad | 170-8841 |
| SYBR® Green Supermix | Bio-Rad | 172-5271 |
| Matrigel |  |  |
| LIF | MedChem | HY-P7084 |
| b-FGF | MedChem | HY-P7004 |
| Mitomycin C | MedChem | HY-13316 |
| Lipofectamine® RNAiMAX |  |  |
| Paraformaldehyde | Thermo Fisher | 30525-89-4 |
| EtOH | Thermo Fisher | H66001GAL |
| Acetone | Thermo Fisher | A18-4 |
| Potassium hydroxide | Thermo Fisher | 1310-28-3 |
| Glycerol | Thermo Fisher | G33-4 |
| Paraffin | Thermo Fisher | 23-021-399 |
| MeOH | Thermo Fisher | BP1105-4 |
| DMSO | Sigma | D2650 |
| H<sub>2</sub>O<sub>2</sub> | Sigma | 216763 |
| Hematoxylin | Sigma | HHS-16 |
| Eosin | Sigma | HT110116 |
| Annexin V | Thermofisher |  |
| TRIzol | ThermoFisher | 15596026 |
| Triton X-100 | Thermo Fisher | BP-151-6 |
| Sodium citrate | Thermo Fisher | S470-212 |
| Hoechst |  |  |
| DNase | NEB | M0303S |
| Cryomatrix | Thermo Fisher | 6769006 |
| **Critical commercial assays** |  |  |
| VECTASTAIN® Universal Quick HRP Kit | Vector<br>Laboratories | Pk-8800 |
| TMR Red for TUNNEL assay | Roche | 12156792910 |
| Qiagen RNeasy Kit | Qiagen | 74104 |
| TURBO DNA-free™ kit | Thermofisher | AM1907 |
| TruSeq Stranded Total RNA Sample Preparation Kit | Illumina | TS-122-2301 |
| iScript cDNA synthesis kit | Bio-rad | 170-8890 |
| ssoAdvanced universal SYBR green supermix | Bio-Rad | 172-5270 |
| **Deposited data** |  |  |
| **Experimental models: Cell lines** |  |  |
| O9-1 cells | Millipore Sigma | SCC049 |
| STO cells | ATCC |  |
| **Experimental models: Organisms/strains** |  |  |
| CD1 | Charles River | Strain code 022 |
| *Wnt-1Cre2* transgenic | Jackson's Lab | 022137 |
| *Gt(ROSA)26Sor<sup>tm4</sup>(ACTB-tdTomato,-EGFP)Luo/J* | Jackson's Lab | 007676 |
| *Trp53<sup>tm1Brn/J</sup>* | Jackson's Lab | 008462 |
| *Sf3b4<sup>L/+</sup>* | Kumar et al<sup>21</sup> | N/A |
| *Sf3b4<sup>L/+</sup>;Wnt1-Cre2<sup>tg/+</sup>* | This study | N/A |
| *Sf3b4<sup>+/-</sup>* | Kumar et al<sup>21</sup> | N/A |
| *Sf3b4<sup>+/-</sup>;Wnt1-Cre2<sup>tg/+</sup>* | This study | N/A |
| *Sf3b4<sup>L/+</sup>;Rosa<sup>mtmg</sup>* | This study | N/A |
| *Sf3b4<sup>L/+</sup>;Trp53<sup>L/+</sup>;Wnt1-Cre2<sup>tg/+</sup>* | This study | N/A |
| *Sf3b4<sup>+/-</sup>;Trp53<sup>L/+</sup>;Wnt1-Cre2<sup>tg/+</sup>* | This study | N/A |
| **Oligonucleotides** |  |  |
| Sf3b4 RNAi | ThermoFisher | 4390771 |
| Negative control RNAi | ThermoFisher | 4390843 |
| Genotyping primers used in this study, see table S5 |  |  |
| qPCR primers used in this study, see table S5 |  |  |
| **Software and algorithms** |  |  |
| ImageJ | NIH | https://imagej.nih.gov/ij/ |
| STAR aligner |  | Version 2.6.1d |
| Gencode M2 |  | Version M2,2013 |
| HTseq |  | Version 0.13.5 |
| rMATS |  | Version 4.0.2 |
| WebLogo |  | Version 3.0 |
| GraphPad Prism 8 Project | GraphPad | Version 8.0.1 |
| **Other** |  |  |
| 24-well plate |  |  |
| Positively charged slides | Thermo Fisher | 1255015 |
| Coverslips | Thermo fisher | 102455 |
| 1.5ml Eppendorf | Progene | 87-B150-C |

## RESOURCE AVAILABILITY

### Lead Contact

Further information and requests for resources and reagents should be directed to and will be fulfilled by the Lead Contact, Loydie A. Jerome-Majewska

### Materials Availability

All antibodies, chemicals and most mouse lines used in this study are commercially available, the details of which can be found in the key resource table and method details section. Details about the mouse lines generated in this study are available from the lead contact upon request.

### Data and Code Availability

RNAseq data have been deposited at [datatype-specific repository] and are publicly available as of the date of publication. Accession numbers are listed in the key resources table.

Microscopy data reported in this paper will be shared by the lead contact upon request. This paper does not report original code.

Any additional information required to reanalyze the data reported in this paper is available from the lead contact upon request.

## EXPERIMENTAL MODELS AND SUBJECT DETAILS

### O9-1 mouse cranial neural crest cell culture

O9-1 cells were purchased from Millipore Sigma (**SCC049**). The cells were cultured on Matrigel (Corning) coated plates in conditioned media supplemented with LIF and b-FGF at final concentrations of 10 ng/ml and 25 ng/ml respectively. Basal O9-1 media was conditioned on mitomycin C inactivated STO cells (purchased from ATCC). LIF (HY-P7084), b-FGF (HY-P7004), and Mitomycin C (HY-13316) were purchased form MedChem Express.

### Transfection of O9-1 cells with siRNAs

O9-1 cells were cultured so that they would be 40% - 60% confluent on the day of transfection. The Lipofectamine® RNAiMAX transfection reagent was used to perform siRNA (10 pmol/μl) transfections. Sf3b4 (4390771) and negative control (4390843) RNAi were purchased from ThermoFisher.

### Mouse lines

All procedures and experiments were performed according to the guidelines of the Canadian Council on Animal Care and approved by the Animal Care Committee of the McGill University Health Centre Research Institute (AUP#5112). Wild type CD1 mice were purchased from Charles Rivers (strain code 022), *Wnt1-Cre2* mice (#022137), the Trp53^tm1Brn^/J mice(#008462) and Gt(ROSA)26Sor^tm4(ACTB-tdTomato,-EGFP)Luo^/J (#007676) were purchased from the Jackson Laboratory.

### Generation of *Sf3b4* mouse line using CRISPR/Cas9

Generation of conditional mutant Sf3b4<em2Lajm> line (*Sf3b4^L/+^*; allele accession ID:MGI:7545559) using CRISPR/Cas9 and Sf3b4<em2.1Lajm> (*Sf3b4^+/-^*; allele accession ID MGI:7545576) was described previously^21^. Briefly, single guide RNAs (sgRNA) were designed in intron 1 and 3 of mouse *Sf3b4.* In the first microinjection using 2gRNAs, a repair template with LoxP sequences and an EcoRV restriction site that targeted exon 3, one male was recovered. He was mated to a wild-type C3H female. One male and one female offspring were confirmed to carry the insertion by Sanger sequencing and they were together to generate homozygous mice with insertion of LoxP site in intron 3. The second round of microinjection, targeting inton1, contained 2gRNAs, a repair template with LoxP sequence and a BamHI site, were carried out in zygotes generated using sperm from a make that was homozygous for LoxP sites in intron 3. Ater identification of pups with integration of LoxP sites in intron 1 and 3, they were mated to C3H wild type mice. One male and one female offspring were verified to have both intron 1 and 3. To establish the *Sf3b4* conditional line (*Sf3b4^L/+^*), one male and one female were backcrossed to the mixed CD1 genetic background or the inbred C57BL/6 genetic background for 10 generations. *Sf3b4*^+/-^ mice were generated by mating *Sf3b4^L^*^/+^ mice on an mixed CD1 genetic background to *β*-*actin* Cre^tg/+^ (*Tmem163^Tg(ACTB-cre^*^) 2Mrt^/CjDswJ; stock #019099, obtained from Jackson Laboratory). Off-springs carrying deletion of exons 2 – 3 were backcrossed to the mixed CD1 genetic background for 10 generation. *Sf3b4^L/+^* and *Sf3b4^+/-^* mice in the mixed CD1 background from 10^th^ generation onwards were used for all mating and analysis in this study.

### Generation of neural crest specific deletion mutation in *Sf3b4*

*Sf3b4^L/+^* and *Sf3b4^+/–^* mice were mated to *Wnt1-cre2^tg/+^* mice to generate *Sf3b4^L/+^;Wnt1-cre2^tg/+^* mice and *Sf3b4^+/–^;Wnt1-cre2^tg/+^* mice respectively. Germline Cre recombinase activity has been previously reported for the *Wnt1-cre2^tg/+^* and was also observed by our lab, therefore no *Sf3b4^L/+^;Wnt1-cre2^tg/+^* male mice were used for this study. *Sf3b4^L/+^;Wnt1-Cre2^tg/+^* mice were mated to *Sf3b4^L/+^* mice, to generate *Sf3b4^L/L^;Wnt1-Cre2^tg/+^* (*Sf3b4^ncc/ncc^*) embryos. Alternatively, *Sf3b4^+/–^;Wnt1-Cre2^tg/+^* mice were mated to *Sf3b4^L/+^* mice to generate embryos with *Sf3b4^L/–^;Wnt1-Cre2^tg/+^* (*Sf3b4^ncc/–^*). *Sf3b4^ncc/–^* embryos generated from both founder lines (male and female; see above) were compared and confirmed to have no difference in phenotype.

## METHOD DETAILS

### Genotyping

Genomic DNA was extracted from mouse tails and embryonic yolk sac using alkaline lysis. To visualize the wildtype, conditional and deletion alleles, a three-primers PCR was used. The following PCR program was done; 30sec 95C, 30sec 60C, 45sec 72C for 28 cycles followed by an elongation step of 5mins at 72C. This PCR amplified the target sequence to generate a wildtype (173bp), a LoxP (213bp) and a mutant (281bp) amplicon. For the commercially available lines, namely *Wnt1-Cre2^tg/+^, R26Sor^mT/mG^* and *Trp53^L/+^* genotyping was conducted as detailed on Jacksons laboratory website: protocol #25394 (*Wnt1-Cre2^tg/+^*), #20368 (*R26Sor^mT/mG^*) and #23419 (*Trp53^L/+^*). See key resources for sequences of all genotyping primers.

### Collection of embryos

Females were placed with a male overnight and checked for vaginal plug in the morning. The day of the plug was considered embryonic day (E)0.5. Dissections were carried out under a Lecia stereomicroscope (Lecia MZ6). Extraembryonic membranes were removed, yolk sac was collected for DNA extraction and genotyping. For embryos collected from E8.5 to E10.5, the pairs of somite was counted. Embryos were collected in 1% or 4% PFA (unless otherwise stated).

### Skeletal preparation

E17.5 embryos we eviscerated and fixed in 100% EtOH overnight. They were then washed into 100%acetone overnight followed by staining with 0.3% Alcian blue/0.1% Alizarin red in acetone for 3-4 days at 37^°^C. The embryos were cleared in 1% potassium hydroxide (KOH) for two days and further cleared using a series of glycerol KOH from 10% to 50%. For long term storage, the skeletal preparations were put in fresh 50% glycerol/1% KOH solution. Skeletal preparations were imaged and analyzed using Lecia stereomicroscope (Lecia MZ6 Infinity1).

### Preparation of Embryos for Embedding and Histology

The dissected embryos were fixed in 4% paraformaldehyde at 4C overnight. For cryo-embedding, the fixed embryos were first cryoprotected in 30% sucrose overnight, embedded in cryomatrix and sectioned sagitally at 10um thickness for immunofluorescence. For embedding in paraffin, fixed embryos were washed in 1xPBS then dehydrated to 100% EtOH, and embedded using paraffin. Embedded embryos were sectioned at 10um thickness on a Lecia RM2155 microtome and mounted on positively charged slides for further staining with hematoxylin and eosin.

### Whole Mount Immunohistochemistry

Embryos were fixed in 4% PFA overnight at 4^0^C, washed and dehydrated into 100% methanol and kept at -20^0^C until use. Embryos were bleached overnight in methanol:DMSO:H_2_O_2_ (4:1:1) solution and then rehydrated into 1xPBS. After several washes with fresh ice-cold PBS+2%milk+5%TritonX-100 (PBSMT), embryos were incubated with 2H3 antibody (DHSB 1:150) overnight. After several short to long washes in PBSMT, embryos were incubated with in HRP conjugated anti-mouse IgG (1:500) overnight. The next day, embryos were washed several time in PBSMT and visualized with diaminobenzidine (DAB, Vector Laboratories). Embryos were visualized and imaged using Leica stereomicroscope (Lecia MZ6 Infinity1)

### Immunofluorescence (IF) and TUNEL Assay

Cryosections of 10um were used for immunofluorescence and TUNEL assay. Immunofluorescence experiments were performed by washing slides in 1XPBS followed by antigen retrieval by heating the slides in 10mM sodium citrate(pH6). After cooling it to room temperature, blocking was carried out in a humidified chamber by covering the slides in 10% normal goat serum in 0.3%tritonX-100/PBS. After 1hr, the blocking solution was drained and replaced with primary antibody: Cleaved Caspase-3 (1:250, Cell Signaling cat#9661T), or αPhospho-histone H3 (Ser10) (1:200, Sigma-Aldrich cat#06-570) and slides were incubated overnight. The next day, slides were washed with PBS, then placed back in the humidified chamber and covered with anti-rabbit Alexa Fluor 568 (1:1000, ThermoFisher Scientific A-11011) secondary antibody. Slides were washed in PBS before mounting. TUNEL assay was carried out using a Cell Death Detection Kit, TMR Red (12156792910, Roche). Slides were fixed in 4% PFA, then washed in PBS, followed by incubation in permeabilization solution in 4C. The slides were washed in PBS then placed in a humidified chamber, covered with TUNEL reaction mix and incubated at 31C for 1hr. Slides were rinsed with PBS, then mounted. Mounting was conducted with Fluroshield aqueous mounting medium with DAPI to visualize the nuclei. Images were captured on a Leica microsystem (model DM6000B) and Leica camera (model DFC 450). For quantification of signal, particle analysis on Image J was used. 3 to 4 sections per embryo were imaged. The percentage of positive cells were determined as follows: the number of PH3 or cleaved Caspase-3-positive cells or TUNEL positive cells/number of DAPI-positive cells (in the pharyngeal arch or hind brain region) X100. Similarly, the percentage of GFP positive cells was determined by: the number of GFP-positive cells/number of DAPI-positive cells X 100. And the percentage of PH3/TUNEL in GFP-positive cells was calculated by: the number of PH3 or TUNEL positive cells/number of GFP-positive cells X 100. These values were plotted using GraphPad (Prism). T-test statistical analysis using the data was also done on GraphPad (Prism)

For O9-1 cells, immunohistochemistry was performed following paraformaldehyde fixation for 20 minutes at room temperature. Cells were washed 3 times with PBS and then permeabolized with 0.1% Triton-X-100/PBS for 15 minutes at room temperature and then blocked with 1% BSA and 5% FBS in 1x PBS for 1 hour. Rabbit anti-PH3 was diluted (1:300) in blocking solution and incubated with cells at room temperature for 1 hour. Cells were then washed 3 times with PBS, followed by incubation with secondary antibodies diluted in blocking solution for 1 hour. DNA was labeled with Hoechst stain. For BrdU, cells were incubated in media supplemented with 10 µM BrdU for 2 h. Cells were then washed and prepared for staining exactly as above with an additional incubation in 2N HCl for 1 h at room temperature after the permeabilization, followed by three PBS washes before continuing the immunostaining with the blocking step. Images were acquired on a Leica SP8 Confocal Laser Scanning microscope. Fluorescence intensity was quantified using Fiji (ImageJ).

### AnnexinV staining of O9-1 cells

O9-1 cells were cultured in 24-well plates so that they were 50%-60% confluent on the day of transfection. *Sf3b4* and negative control siRNAs were transfected using Lipofectamine® RNAiMAX as described above. 24 hours after transfection, the Annexin V (ThermoFisher) staining solution was added to the wells (500 μl for a 24-well plate) and incubated, in the dark, for 15 minutes. The Annexin V staining solution was aspirated form the wells and the cells were washed once with 1X Annexin V binding buffer and then quantified on a microplate reader.

### Reactive Oxygen Species (ROS) and Superoxide staining of O9-1 cells

O9-1 cells were seeded in a 96-well black wall/clear bottom plate and transfected with *Sf3b4* and negative control siRNAs at 50% - 60% confluency. 24 hours later, the Oxidative Stress Detection Reagent and the Superoxide Detection Reagent (Abcam) were added and incubated with the cells for 1 hour. Quantification was performed in a microplate reader set to standard fluorescein (Excitation = 488 nm and Emission = 520 nm). Five readings (technical replicates) were taken. The filter set was then changed to rhodamine (Excitation = 550 nm and Emission = 610 nm) and another five reading were taken.

### RNA isolation of embryos

For RNA isolation, head from E8.5 embryos of 8-10 somites were collected in 250µl of RNAlater (Invitrogen AM7020). For each pool of RNA extraction, head from 3 embryos were pooled. RNA extraction was carried out using the Qiagen RNeasy micro kit per manufacturer’s instruction (Qiagen 74104). For RNA sequencing, three wild type and three mutant pools were generated with each pool containing heads from 3-embryos.

### RT-PCR and qPCR using RNA from embryos

Total RNA was treated with DNAse I (NEB, M0303S) according to manufacturer’s protocol and used for reverse transcription with iScript cDNA synthesis kit (Bio-rad, cat #170-8890) according to the manufacturer’s protocol. The RT-PCR program included a hot start at 95°C for 5 min, followed by 40 cycles of denaturation at 95°C for 10s and annealing/extension at 60°C for 30s and elongation for 30s. Each experiment was performed twice, using five pools of wildtype, and five *Sf3b4^ncc/-^* RNA. qPCR experiments were performed using the ssoAdvanced universal SYBR green supermix (Bio-Rad, cat#172–5270) on a Roche LightCycle 480 PCR machine, these were done in duplicates to ensure technical replicability and target genes were normalized using GAPDH. Sequences for all primers used are listed in Key resources.

### RNA isolation and qPCR of O9-1 cells

RNA was collected in 500 μl of TRIzol reagent (cat# 15596026, Thermo Fischer Scientific) 36 hours post-transfection. RNA was extracted following the manufacturer’s protocol and recommendations. The TURBO DNA-free™ kit (AM1907, Thermo Fisher Scientific) was used to digest any DNA that might have been present in the RNA sample. cDNA was generated from RNA at a concentration of 400ng with iScript RT Supermix for RT-qPCR (BioRad). Expression levels were calculated relative to the average expression of 3 housekeeping genes via the ΔΔCt method.

### RNA sequencing Analysis

Sequencing libraries were prepared by the McGill Genome Centre (Montreal, Canada), using the TruSeq Stranded Total RNA Sample Preparation Kit (Illumina TS-122-2301) by depleting ribosomal and fragmented RNA, synthesizing first- and second strand complementary DNA (cDNA), adenylating the 3′ ends and ligating adaptors, and enriching the adaptor-containing cDNA strands by PCR. The libraries were sequenced using an Illumina NovaSeq 6000 PE100sequencer, with 100 nucleotide paired-end reads, generating between 109and 230 million reads sample. The sequencing reads were trimmed using CutAdapt (Martin, 2011) and mapped to the mouse reference genome(mm10) using STAR (Dobin et al., 2013) aligner (version 2.6.1d), with default parameters, and annotated using the Gencode (Harrow et al., 2006) M2 (version M2, 2013) annotation. Htseq-count [ part of the ‘HTSeq’(Anders et al., 2015) framework, version 0.13.5] was used for expression quantification.

Differential expression analysis was done as using the DESeq2^58^ package and a list of significant DEGs was derived using an FDR cutoff of less than 0.05 with no additional restriction on the absolute log2 fold change (Log2FC) (to allow for detection of even minor expression changes).

A comprehensive Overrepresentation Analysis (ORA) to gain insights into the biological processes, molecular functions, cellular components, and KEGG pathways associated with our gene clusters was conducted. The analysis was performed using the clusterProfiler^59^, enrichplot^60^, and org.Mm.eg.db^61^ R packages, which are widely recognized for their robust capabilities in functional enrichment analysis for genomic data.

For Gene Ontology Biological Process (GO BP), we identified significantly enriched terms that highlight the key biological activities associated with our gene clusters.

The clusterProfiler package facilitated the statistical analysis of functional profiles, enrichplot was instrumental in visualizing the enriched terms in an interpretable manner, and org.Mm.eg.db ensured accurate gene annotation and mapping. The integration of these packages enabled a robust interpretation of the biological significance of our gene clusters.

RMATS 4.1.2^62^ was used for differential splicing analysis. Detected splicing events were filtered by systematically excluding those with a mean of inclusion junction counts (IJC) lower than 5 in either wild-type or mutant samples. For identifying significant differential spliced events (DSE) and absolute inclusion level difference (ILD), we used a cut-off of more than 0.05 and a Benjamin-Hochberg multiple testing correction with a false discovery rate (FDR) cut-off of less than 0.05. The FDR cut-off was relaxed to obtain a large dataset enriched for alternative splicing (AS) events in order to observe general tendencies, such as increased propensity for exon skipping or intron retention. To characterize 3’SS sequences, LaBranchoR^63^, a branchpoint (BP) prediction tool was used. The BPs and their surrounding area consensus motifs were generated using WebLogo 3.0^64^.

Enrichment analysis using the Mouse Genome Informatics (MGI) for mammalian phenotype was done using skipped exon data that was mutant specific on events with p-value of <0.05, ILD of >0.05 and FDR of <0.05.

## QUANTIFICATION AND STATISTICAL ANALYSIS

Details for data analysis and software used are detailed in each method section. No additional statistical analyses were conducted. Figure legends contain number of replicates and details of data plotted (mean, SEM).

## Notes

### Competing Interest Statement

The authors have declared no competing interest.

