## supplemental figure and tables for "Etiology of Craniofacial and Cardiac Malformations in a Mouse Model of SF3B4-Related Syndromes"

Figure S1

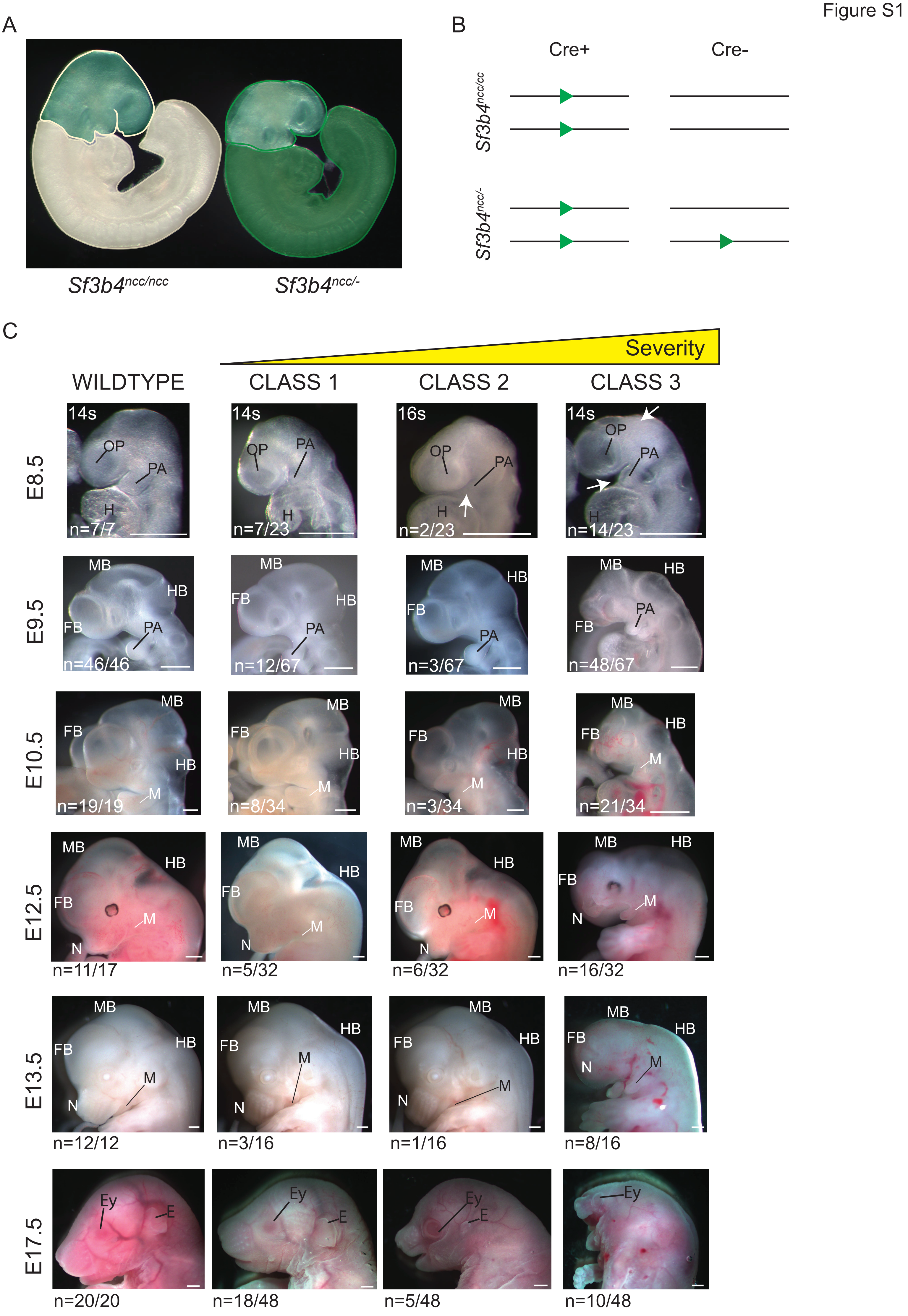
A
Sf3b4ncc/ncc
Sf3b4ncc/-
B
Cre+
Cre-
Sf3b4ncc/cc
Sf3b4ncc/-
C
Severity
WILDTYPE
CLASS 1
CLASS 2
CLASS 3
E8.5
14s
OP
PA
H
n=7/7
14s
OP
PA
H
n=7/23
16s
OP
PA
H
n=2/23
14s
OP
PA
H
n=14/23
E9.5
MB
HB
FB
PA
n=46/46
n=12/67
n=3/67
n=48/67
E10.5
MB
FB
HB
M
n=19/19
n=8/34
n=3/34
n=21/34
E12.5
MB
HB
FB
M
N
n=11/17
n=5/32
n=6/32
n=16/32
E13.5
MB
FB
HB
M
N
n=12/12
n=3/16
n=1/16
n=8/16
E17.5
Ey
E
n=20/20
n=18/48
n=5/48
n=10/48

**Figure S1: Neural crest specific mutations in *Sf3b4* in mouse embryos lead to craniofacial defects with variable expressivity**

A. Schematic to show the two different mutants used to study the loss of *Sf3b4* during craniofacial development. *Sf3b4*$^{ncc/ncc}$ have homozygous deletion of *Sf3b4* in NCCs (green) and wild type *Sf3b4* in neighbouring cell (beige). *Sf3b4*$^{ncc/-}$ embryos have homozygous deletion of *Sf3b4* in NCCs while neighbouring cells have heterozygous deletion of *Sf3b4* (dark green).
B. Schematic to show the *Sf3b4* alleles in cells that are CRE+ and CRE-. *Sf3b4*$^{ncc/ncc}$ have homozygous deletion of *Sf3b4* in CRE+ cells and wild type *Sf3b4* in neighbouring CRE- cells. *Sf3b4*$^{ncc/-}$ embryos have homozygous deletion of *Sf3b4* in NCCs while neighbouring CRE- cells have heterozygous deletion of *Sf3b4*.
C. Representative images of embryos at E8.5-17.5 showing variable expressivity of craniofacial abnormalities observed in *Sf3b4* NCC mutant embryos compared to wildtype. Embryos images of Class 1 and 2 are *Sf3b4*$^{ncc/ncc}$. Class 1 embryos at all stages are comparable to wildtype. At E8.5 embryos 14-16s are shown. At this stage Class 2 embryo has a hypoplastic 1$^{st}$ PA. From E9.5-E13.5, Class 2 embryos have a normal craniofacial region with hypoplastic mid- and hind-brain. All Class 3 embryo images are *Sf3b4*$^{ncc/-}$. Class 3 embryos are severely affected with hypoplastic PA as well as hypoplastic fore-, mid- and hind-brain at early stages. These embryos have unfused frontonasal prominence at E10.5 and absence of nasal region from E12.5 onwards. Otic vesicle/external ears and eyes are absent at later stages along with an exposed forebrain. At E8.5 the white arrows point to hypoplastic structures. FB: forebrain, FL: forelimb, H: heart, HB: hindbrain, HL: hindlimb, M:mandible, MB:midbrain, N:nasal, OP: optic placode, PA: Pharyngeal arch. E8.5-13.5, scale bar is 500μm. E17.5, scale bar is 1000μm.

Figure S2

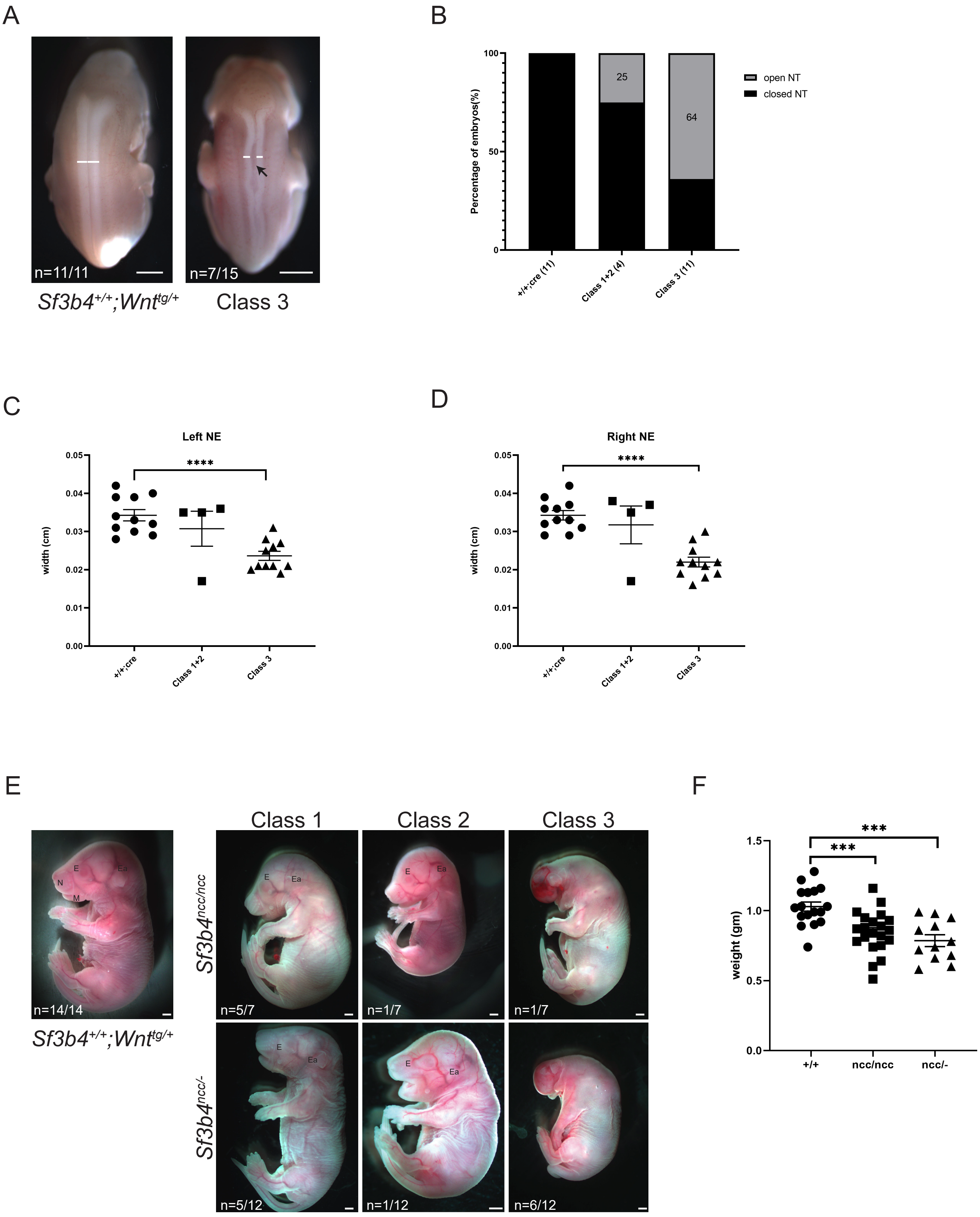
A
n=11/11
n=7/15
Sf3b4+/+;Wnttg/+
Class 3
B
Percentage of embryos(%)
100
50
0
25
64
open NT
closed NT
+/+;cre (11)
Class 1+2 (4)
Class 3 (11)
C
Left NE
****
width (cm)
0.05
0.04
0.03
0.02
0.01
0.00
+/+;cre
Class 1+2
Class 3
D
Right NE
****
width (cm)
0.05
0.04
0.03
0.02
0.01
0.00
+/+;cre
Class 1+2
Class 3
E
Class 1
Class 2
Class 3
N
E
Ea
M
n=14/14
Sf3b4+/+;Wnttg/+
Sf3b4ncc/ncc
n=5/7
n=1/7
n=1/7
Sf3b4ncc/-
n=5/12
n=1/12
n=6/12
F
***
***
weight (gm)
1.5
1.0
0.5
0.0
+/+
ncc/ncc
ncc/-

**Figure S2: Open neural tube was observed in several *Sf3b4* mutant embryos at E12.5**

A. Representative images at E12.5 of wild type embryos and open neural tube in Class 3 mutant embryos. Dashed white line shows where measurement for the width of the neural tube was taken – in line with the anterior most part of the forelimb. Black arrow indicates open neural tube. Scale bar is 1000µm
B. Graphical representation of percentage of mutant embryos in each Class that have open neural tube.
C. Graphical representation showing significant decrease in the width of the left neural epthelium in mutant embryos. Each data point represents one embryo. *$p<0.05$, ****$p<0.0001$.
D. Graphical representation showing significant decrease in the width of the right neural epithelium in mutant embryos. Each data point represents one embryo. *$p<0.05$, ****$p<0.0001$.
E. Representative images of E17.5 wildtype as well as mutant embryos with Class 1, Class 2 and Class 3 that were used for skeletal preparation.
F. Graph showing significant decrease in weight of E17.5 *Sf3b4*$^{ncc/ncc}$ and *Sf3b4*$^{ncc/-}$ embryos (*Sf3b3*$^{+/+}$ n=17, *Sf3b4*$^{ncc/ncc}$ n=21, *Sf3b4*$^{ncc/-}$ n=12). Each data point indicates one embryo and error bars are ± s.e.m, ***$p<0.001$.

Figure S3

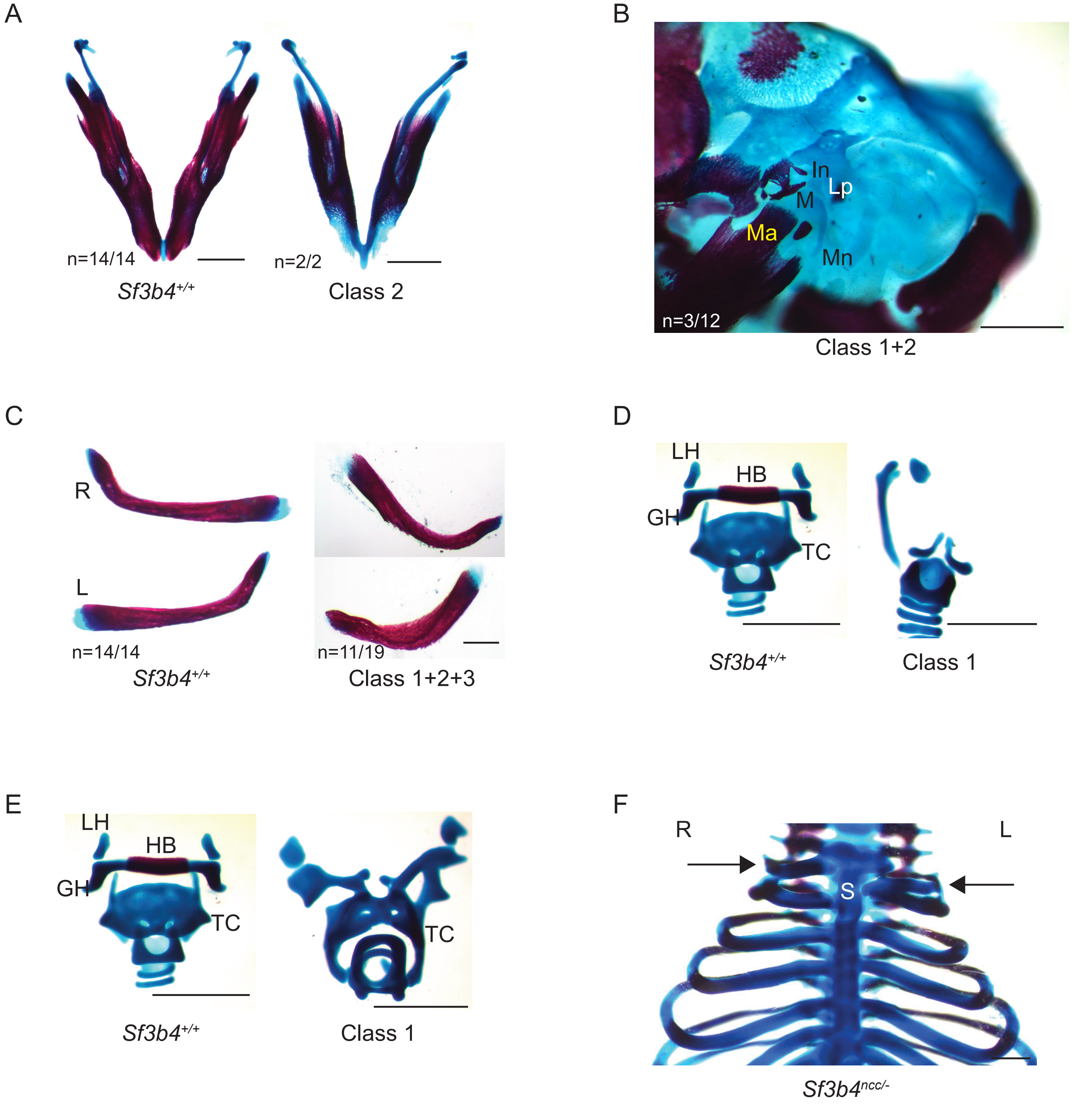
A
n=14/14
Sf3b4+/+
n=2/2
Class 2
B
In
Lp
M
Ma
Mn
n=3/12
Class 1+2
C
R
L
n=14/14
Sf3b4+/+
n=11/19
Class 1+2+3
D
LH
HB
GH
TC
Sf3b4+/+
Class 1
E
LH
HB
GH
TC
Sf3b4+/+
TC
Class 1
F
R
L
S
Sf3b4ncc/-

**Figure S3: Skeletal defects are observed in craniofacial and axial regions in *Sf3b4* mutants, related to Figure 1.**

A. Representative image showing decreased ossification in mandible of Class 2 mutant compared to wild type. Image is of *Sf3b4*$^{ncc/ncc}$. Sale bar is 1000μm.
B. Representative image of hypoplastic middle ear structures observed in Class1 and 2 *Sf3b4* mutant embryos. Image is of *Sf3b4*$^{ncc/ncc}$. In: incus, Lp: lenticular process, M: malleus, Ma: mandible, Mn: manubrium. Scale bar is 1000μm.
C. Representative image of hypoplastic and curved clavicle in *Sf3b4* NCC mutants of all Classes compared to wild type. L: left clavicle, R: right calvicle. Scale bar is 100μm.
D. Representative image of vertically placed hypoplastic hypoid body with no ossification, and absent thyroid cartilage in one Class 1 mutant. GH: greater horn, HB: hyoid body, LB: lesser horn, TC: thyroid cartilage. Scale bar is 1000μm.
E. Representative image of malformed hypoid body and thyroid cartilage in one Class 1 mutant. TC: thyroid cartilage. Scale bar is 1000μm
F. Representative image of ectopic rib found at C7 in *Sf3b4*$^{ncc/-}$ mutant embryos. Arrow shows the rib arising from C7, attached to the sternum on the right side and attached to the T1 rib on the left side. L: left, R: right, S: sternum. Scale bar is 1000μm.

Figure S4

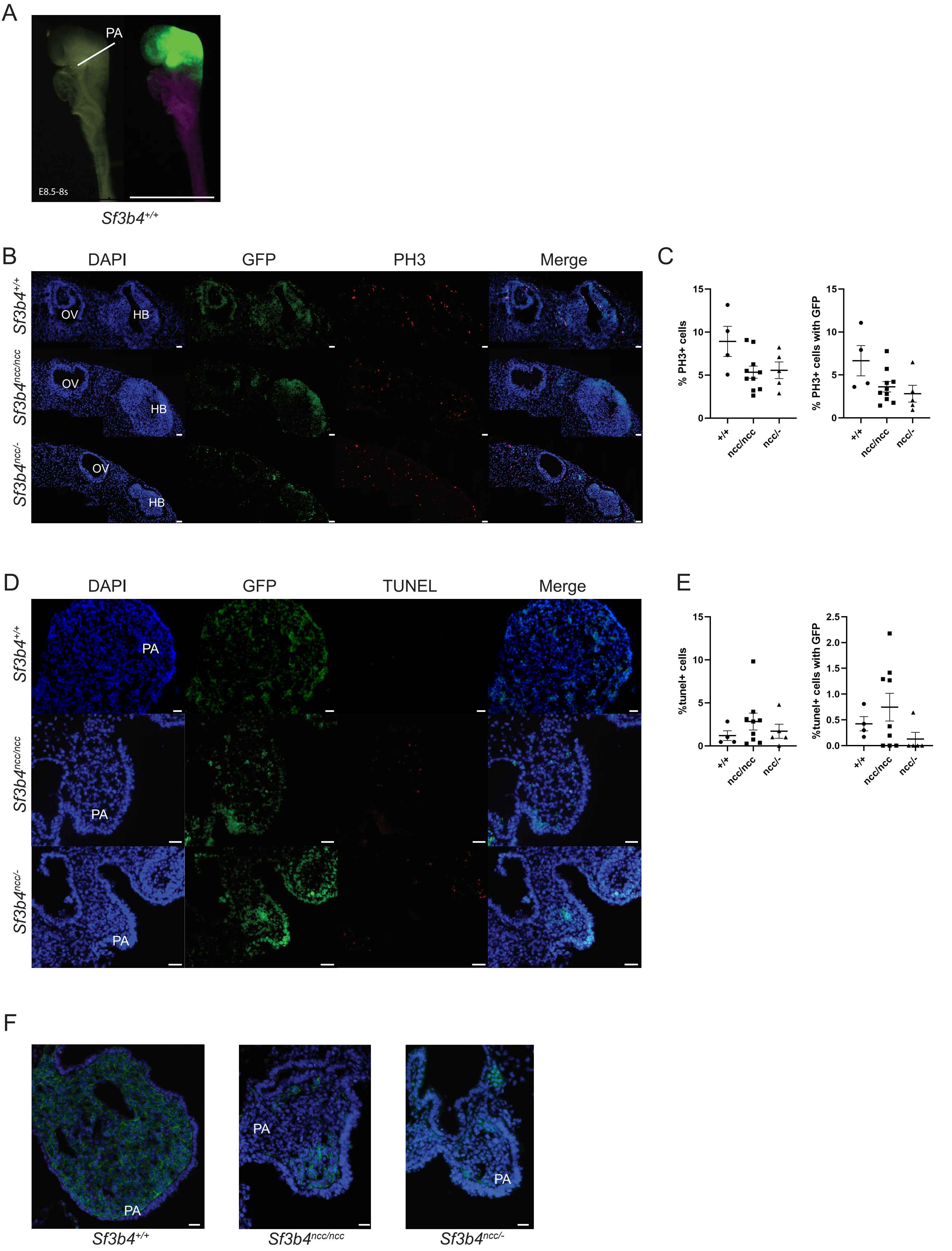

A
PA
E8.5-8s
Sf3b4+/+
B
DAPI
GFP
PH3
Merge
Sf3b4+/+
Sf3b4ncc/ncc
Sf3b4ncc/-
OV
HB
C
% PH3+ cells
% PH3+ cells with GFP
+/+
ncc/ncc
ncc/-
D
DAPI
GFP
TUNEL
Merge
PA
E
%tunel+ cells
%tunel+ cells with GFP
F
Sf3b4+/+
Sf3b4ncc/ncc
Sf3b4ncc/-

**Figure S4: NCCs with mutation in *Sf3b4* have decreased proliferation and increased cell death, related to Figure 4.**

A. E8.5-8s wild type embryo with GFP expression in the head and $1^{st}$ pharyngeal arch. PA; pharyngeal arch. Scale bar is 20μm.
B. Representative images of PH3 staining on sections with hind brain and otic vesicle region of wild type and mutant embryos to look at proliferation. OV; otic vesicle, HB; hindbrain. Scale bar is 20μm.
C. Graph showing quantification of PH3+ cells in the hindbrain and otic vesicle region. Each data point represents the average of 4 sections from one embryo, error bars are ±s.e.m.
D. Representative images of TUNEL assay on sections of the $1^{st}$ pharyngeal arch in wild type and mutant embryos. PA; pharyngeal arch. Scale bar is 20μm.
E. Graph showing quantification of TUNEL+ cells in the pharyngeal arch. Each data point represents the average of 3/4 sections from one embryo, error bars are ±s.e.m.
F. Representative images of caspase-3 staining on sections of the $1^{st}$ pharyngeal arch in wild type and mutant embryos showing no caspase-3 in all. Scale bar is 20μm.

Figure S5

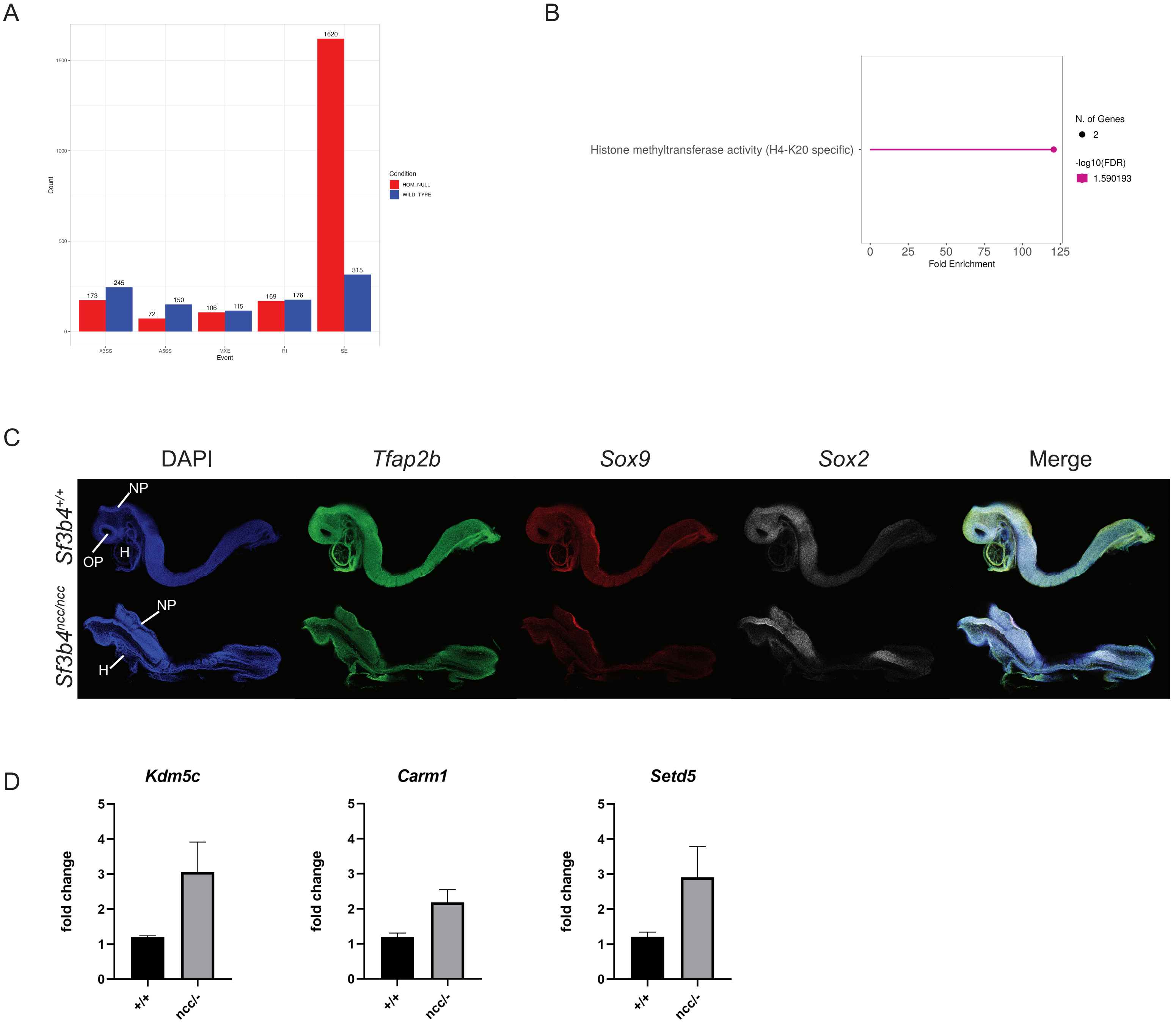
A
B
C
D
1620
1500
1000
500
0
Count
173
245
72
150
106
115
169
176
315
A3SS
A5SS
MXE
RI
SE
Event
Condition
HOM_NULL
WILD_TYPE
Histone methyltransferase activity (H4-K20 specific)
N. of Genes
2
-log10(FDR)
1.590193
0
25
50
75
100
125
Fold Enrichment
DAPI
Tfap2b
Sox9
Sox2
Merge
Sf3b4+/+
Sf3b4ncc/ncc
NP
OP
H
Kdm5c
Carm1
Setd5
fold change
5
4
3
2
1
0
+/+
ncc/-

**Figure S5: Significant changes in differentially expressed genes and differentially skipped event is observed by RNAseq, related to Figure 5.**

A. Graph showing number of events that were differentially spliced for each of the splicing events.
B. Gene ontology for A5SS events.
C. HCR of *Tfap2b*, *Sox 9* and *Sox2* in wild type (n=3) and *Sf3b4*$^{ncc/ncc}$ (n=3) embryos showing decreased expression of *Tfap2b* and *Sox9* in mutants. H; heart, NP; neural plate, OP; otic placode.
D. RT-qPCR showing fold change of *Kdm5c*, *Carm1* and *Setd5* relative to *Gapdh* in wild type (n=3) and mutants (n=3).

Figure S6

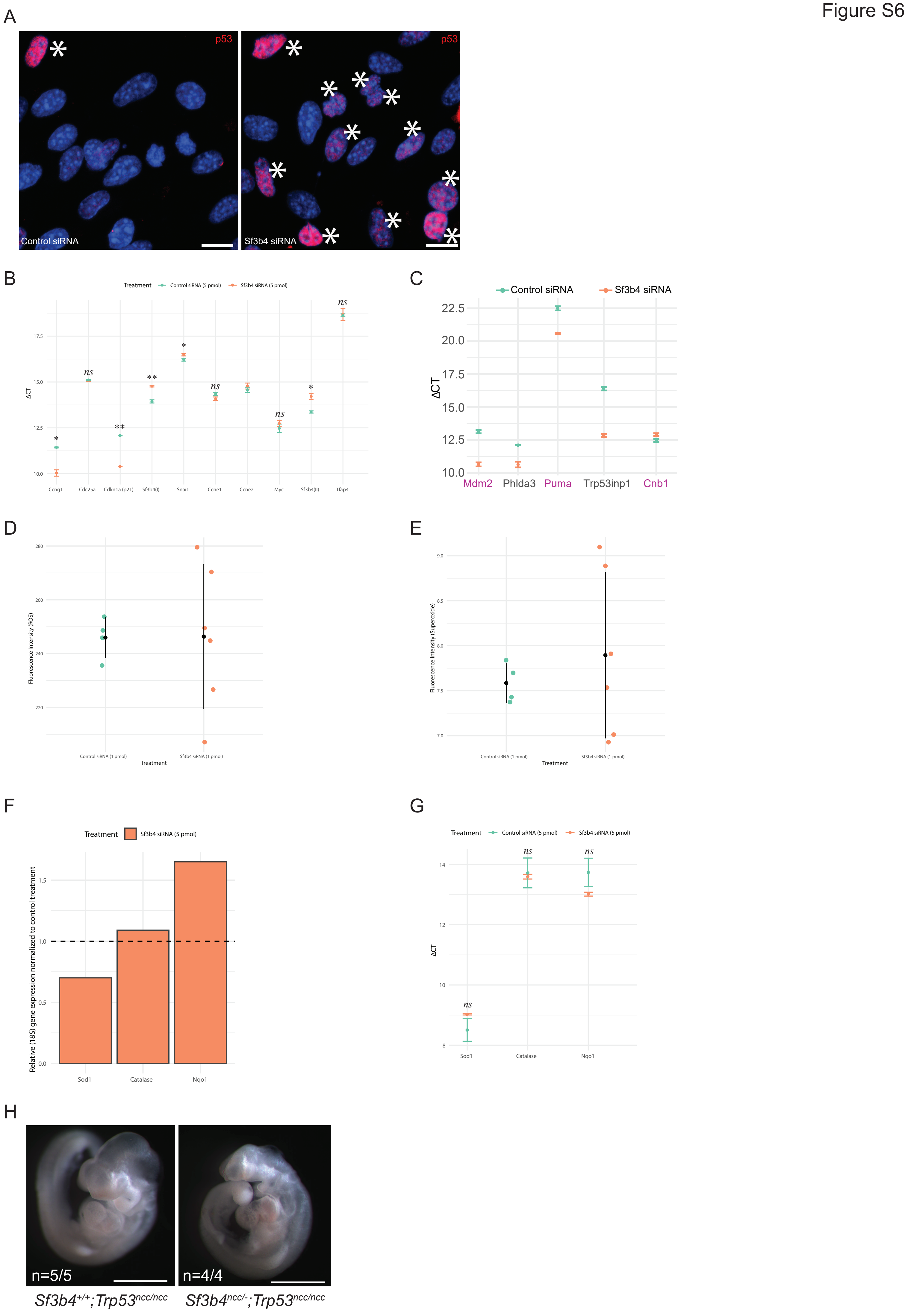
A
p53
Control siRNA
p53
Sf3b4 siRNA
B
Treatment
Control siRNA (5 pmol)
Sf3b4 siRNA (5 pmol)
ΔCT
17.5
15.0
12.5
10.0
ns
*
**
ns
ns
*
ns
*
**
ns
Ccng1
Cdc25a
Cdkn1a (p21)
Sf3b4(I)
Snai1
Ccne1
Ccne2
Myc
Sf3b4(II)
Tfap4
C
Control siRNA
Sf3b4 siRNA
ΔCT
22.5
20.0
17.5
15.0
12.5
10.0
Mdm2
Phlda3
Puma
Trp53inp1
Cnb1
D
Fluorescence Intensity (ROS)
280
260
240
220
Control siRNA (1 pmol)
Sf3b4 siRNA (1 pmol)
Treatment
E
Fluorescence Intensity (Superoxide)
9.0
8.5
8.0
7.5
7.0
Control siRNA (1 pmol)
Sf3b4 siRNA (1 pmol)
Treatment
F
Treatment
Sf3b4 siRNA (5 pmol)
Relative (18S) gene expression normalized to control treatment
1.5
1.0
0.5
0.0
Sod1
Catalase
Nqo1
G
Treatment
Control siRNA (5 pmol)
Sf3b4 siRNA (5 pmol)
ΔCT
14
12
10
8
ns
ns
ns
Sod1
Catalase
Nqo1
H
n=5/5
n=4/4
Sf3b4+/+;Trp53ncc/ncc
Sf3b4ncc/-;Trp53ncc/ncc

**Figure S6: NCCs with *Sf3b4* mutation have increased P53 activity.**

A. Representative images of control or Sf3b4 sRNA treated O9-1 cells with p53 staining. White * indicates cells where p53 is observed. Scale bar is 10µm.
B. Graph showing change in expression levels of cell cycle genes in the p53 pathway, the ΔCT values of O9-1cells treated with control and Sf3b4 siRNA is plotted.
C. Graph showing change in expression levels of genes in the p53 apoptosis pathway, the ΔCT values of O9-1cells treated with control and Sf3b4 siRNA is plotted.
D. ROS levels in Sf3b4 knockdown O9-1 cells (n=4) compared to control (n=6). A strip chart of fluorescence intensity, in arbitrary units, associated with ROS (excitation λ = 488 nm, emission λ = 520 nm) for control (green) and knockdown (orange) conditions. (B) Strip chart of fluorescence intensity, in arbitrary units, associated with Superoxide (excitation λ = 550 nm, emission λ = 610 nm) for control (green) and knockdown (orange) conditions. Black dot represents data mean and error bars are standard deviations.
E. Superoxide levels in Sf3b4 knockdown O9-1 cells (n=4) compared to control (n=6). A strip chart of fluorescence intensity, in arbitrary units, associated with Superoxide (excitation λ = 550 nm, emission λ = 610 nm) for control (green) and knockdown (orange) conditions. Black dot represents data mean and error bars are standard deviations.
F. Expression of Sod1, Catalase, and Nqo1 in Sf3b4 knockdown O9-1 cells relative to control presented, as average fold-change. Horizontal dashed line represents expression of the abovementioned genes in the control cells.
G. Average ΔCT values of each gene for knockdown (orange) and control (green) samples. Error bars represent standard deviations (n=2). The significance of the difference between the means was tested using a one-tailed, heteroscedastic, Student’s t-test (ns p>0.05).
H. Representative images of E9.5 wild type and *Sf3b4* NCC mutant embryos with decreased levels of p53, showing no rescue of the craniofacial phenotype in mutants. Scale bar is 1000µm.

### Figure S7

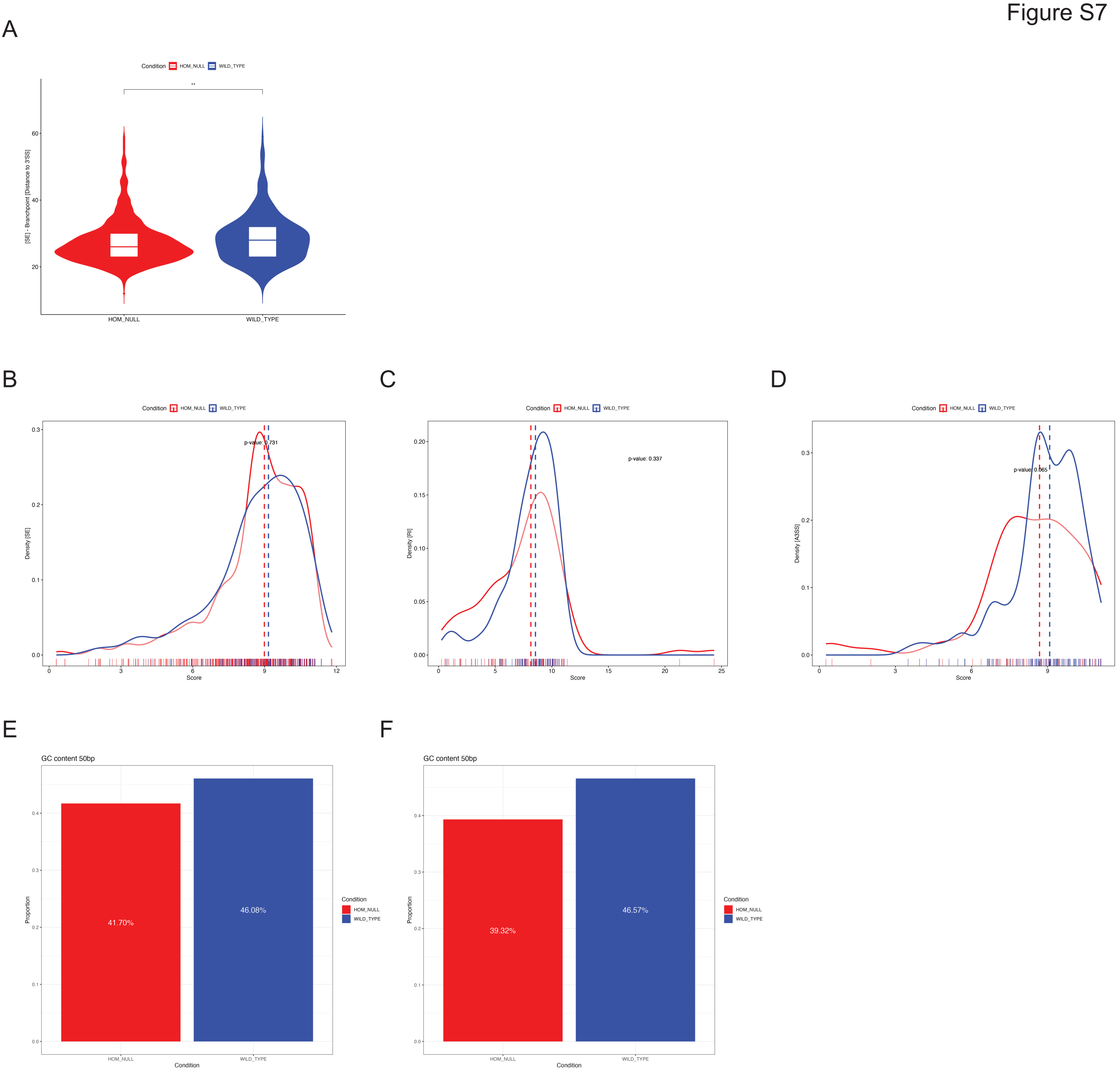

A
Condition HOM_NULL WILD_TYPE
**
[SE] - Branchpoint [Distance to 3'SS]
20
40
60
HOM_NULL
WILD_TYPE
B
Condition HOM_NULL WILD_TYPE
p-value: 0.731
Density [SE]
Score
C
Condition HOM_NULL WILD_TYPE
p-value: 0.337
Density [RI]
Score
D
Condition HOM_NULL WILD_TYPE
p-value: 0.065
Density [A3SS]
Score
E
GC content 50bp
Proportion
41.70%
46.08%
Condition
HOM_NULL
WILD_TYPE
F
GC content 50bp
Proportion
39.32%
46.57%
Condition
HOM_NULL
WILD_TYPE

**Figure S7: Branch point analysis of alternatively spliced events, related to Figure 5.**

A. Violin plot showing predicted branch point distances from 3’SS in wild type and mutants. **p<0.01.

B-D. Graphs showing weaker predicted 5’SS strength of SE, RI and A5SS events respectively.

E-F. Graphs showing decreased GC content in SE and MXE events respectively.

Supplementary Table 1.1: Percentage of *Sf3b4*$^{ncc/ncc}$ embryos that were abnormal and reabsorbed at each stage. Number with 'L' refers to the number of litters examined at each stage

| *Sf3b4*$^{ncc/ncc}$ | E8.5(17L,63) | E9.5(14L,22) | E10.5(6L,14) | E12.5(10L,21) | E13.5(3L,7) | E17.5(12L,32) |
|---|---|---|---|---|---|---|
| %Abnormal | 16.9 | 59.1 | 42.9 | 61.9 | 71.4 | 28.1 |
| %Reabsorbed | 0.0 | 0.0 | 0.0 | 14.3 | 14.3 | 34.4 |
| Total% | 16.9 | 59.1 | 42.9 | 76.2 | 85.7 | 62.5 |

Supplementary Table 1.2: Percentage of *Sf3b4*$^{ncc/-}$ embryos that were abnormal and reabsorbed at each stage.

| *Sf3b4*$^{ncc/-}$ | E8.5(80L,168) | E9.5(20L,45) | E10.5(7L,20) | E12.5(4L,11) | E13.5 (6L,9) | E14.5(5L,6) | E17.5(9L,16) |
|---|---|---|---|---|---|---|---|
| %Abnormal | 18.5 | 80.0 | 85.0 | 63.6 | 66.7 | 33.3 | 50.0 |
| %Reabsorbed | 0.6 | 2.2 | 5.0 | 27.3 | 33.3 | 66.7 | 18.8 |
| Total% | 19.1 | 82.2 | 90.0 | 100 | 100 | 100 | 68.8 |

Supplementary Table 2.1: Distribution of each genotype from *Sf3b4*$^{L/+}$;*Wnt1-Cre2*$^{tg/+}$ and *Sf3b4*$^{L/L}$ mating and chi-square analysis showing significant loss of *Sf3b4*$^{ncc/ncc}$ pups

| Age of pup | Genotype | | | |
|---|---|---|---|---|
| | L/+ | L/L | L/+;Wntcre/+ | L/L;wntcre/+ |
| P0 (1L, 6) | 2 | 1 | 2 | 0 |
| P21 (6L, 62) | 20 | 21 | 20 | 2 |
| Total (7L, 68) | 22 | 22 | 22 | 2 |
| $\chi2$=17.64, p=0.0052 | | | | |

Supplementary Table 2.2: Distribution of each genotype from *Sf3b4*$^{+/-}$;*Wnt1-Cre2*$^{tg/+}$ and *Sf3b4*$^{L/+}$ mating and chi-square analysis showing significant loss of *Sf3b4*$^{ncc/-}$ pups

| Age of pup | Genotype | | | | | | | |
|---|---|---|---|---|---|---|---|---|
| | +/+ | +/- | L/+ | L/- | +/+;wntcre/+ | +/-;wntcre/+ | L/+;wntcre/+ | L/-;wntcre/+ |
| P21(2L, 13) | 5 | 1 | 1 | 4 | 1 | 0 | 1 | 0 |
| $\chi2$=14.69, p=0.065 | | | | | | | | |

Supplementary Table 3: Number of E17.5 *Sf3b4* NCC male and female mutants observed at each Class

| Phenotype | Male (21) | Female (17) |
|---|---|---|
| Class 1 | 5 | 7 |
| Class 2 | 2 | 4 |
| Class 3 | 7 | 2 |
| Reabsorbed | 7 | 4 |

Supplementary Table 4: Number and percentage of embryos with specific skeletal abnormalities observed at E17.5 in *Sf3b4* NCC mutant embryos and their tissue of origin. *Abnormalities in these structures were only observed in *Sf3b4*$^{ncc/-}$ embryos

| **Structure** | **Defect** | **Number of embryos (19)** | **%** | **Tissue of origin** |
|---|---|---|---|---|
| ***Cranium*** | | | | |
| Frontal | Decreased mineralization | 3 | 16 | NCC |
| | Cartilage | 3 | 16 | |
| | Absent | 7 | 37 | |
| Parietal | Decreased mineralization | 3 | 16 | Mesoderm |
| | cartilage | 4 | 21 | |
| | Absent | 7 | 37 | |
| Inter-parietal | cartilage | 5 | 26 | NCC+Mesoderm |
| Supraoccipital | cartilage | 9 | 47 | Mesoderm |
| Basisphenoid | Malformed | 12 | 63 | NCC |
| Pterygoid | Small | 2 | 11 | NCC |
| | Malformed | 3 | 16 | |
| | Absent | 8 | 42 | |
| ***Palatal*** | | | | |
| Cleft | | 6 | 32 | |
| Palatal process of maxilla | Malformed | 2 | 11 | NCC |
| | Absent | 7 | 37 | |
| Palatal process of premaxilla | Absent | 7 | 37 | NCC |
| Premaxilla | Absent | 7 | 37 | NCC |
| Palatal process of palatine | Absent | 10 | 53 | NCC |
| Palatine | Malformed | 2 | 11 | NCC |
| | Absent | 8 | 42 | |
| ***Mandible*** | | | | |
| Meckle’s Cartilage | Small | 1 | 5 | NCC |
| | Malformed | 1 | 5 | |

| | | | | |
|---|---|---|---|---|
| | Absent | 7 | 37 | |
| Mandible | Small | 3 | 16 | NCC |
| | Malformed/absent | 9 | 47 | |
| ***Middle ear*** | | | | |
| Malleus | Malformed | 3 | 16 | NCC |
| | Absent | 7 | 37 | |
| Incus | Malformed | 3 | 16 | NCC |
| | Absent | 7 | 37 | |
| Stapes | Absent | 13 | 68 | NCC |
| Tympanic ring | Small | 1 | 5 | NCC |
| | Absent | 10 | 53 | |
| Styloid process | Malformed | 1 | 5 | NCC |
| | Absent | 10 | 53 | |
| ***Others*** | | | | |
| Hyoid | Malformed | 2 | 11 | NCC |
| | Absent | 10 | 53 | |
| Thyroid cartilage | Malformed | 1 | 5 | NCC |
| | Absent | 9 | 47 | |
| Clavicle | Malformed | 11 | 58 | NCC |
| Ribs | Ectopic rib at C7 | 7 | 37 | Mesoderm |
| | Unevenly spaced ribs | 1 | 5 | Mesoderm |
| | $8^{th}$ rib attached to sternum | 1 | 5 | Mesoderm |
| Vertebrae* | T1 to T2 | 1 | 5 | Mesoderm |
| | Missing L6 | 10 | 53 | Mesoderm |
| | Iliac attached to S2 | 2 | 11 | Mesoderm |
| | Open neural tube | 4 | 21 | Mesoderm |
| | Curved spine | 3 | 16 | Mesoderm |

Supplementary Table 5: Significantly DEGs involved in neural crest development and their fold change.

| **Gene** | **Fold change** | **padj-value** |
|---|---|---|
| *Tfap2b* | 0.504 | 1.49E-10 |
| *Sox5* | 0.673 | 2.93E-4 |
| *Sox6* | 0.756 | 3.02E-2 |
| *Sox9* | 0.811 | 4.22e-2 |
| *Sox10* | 0.489 | 2.14E-13 |
| *Ets1* | 0.841 | 3.49E-2 |
| *En2* | 0.417 | 2.25E-07 |
| *Dnmt1* | 0.848 | 3.22E-2 |
| *Dnmt3b* | 0.849 | 4.90E-2 |
| *Nes* | 0.731 | 1.02E-5 |
| *Grhl3* | 1.616 | 1.54E-3 |

| | | |
|---|---|---|
| *E-cad* | 1.255 | 2.69E-2 |

Supplemental Table 6: List of primers used in this manuscript and their sequences.

| Primer name | Sequence |
|---|---|
| **Genotyping primers** | |
| Sf3b4 genotyping F | CCCGGAAGTGGAAGTTGTAA |
| Sf3b4 genotyping R | AACGCTTCCGGCACTAATCT |
| Sf3b4 genotyping R2 | ACTCCATCTGCCAACTTCATC |
| Wnt Transgene F | CAGCGCCGCAACTATAAGAG |
| Wnt Transgene R | CATCGACCGGTAATGCAG |
| Wnt Internal Positive Control | CAAATGTTGCTTGTCTGGTG |
| Wnt Internal Positive Control | GTCAGTCGAGTGCACAGTTT |
| Trp53 F | GGTTAAACCCAGCTTGACCA |
| Trp53 R | GGAGGCAGAGACAGTTGGAG |
| Rosa mT/mG Common | CTTTAAGCCTGCCCAGAAGA |
| Rosa mT/mG Mutant F | TAGAGCTTGCGGAACCCTTC |
| Rosa mT/mG Wild type F | AGGGAGCTGCAGTGGAGTAG |
| **RT-PCR primers** | |
| Tcf7l2 F | CCCCTTCATCATGATCCCCG |
| Tcf7l2 R | GTTTGTCTGCTCTGGAGGCT |
| Kdm5c F | TTCAGATGGTGGAGAGGGCA |
| Kdm5c R | GTCACCTCAAGCAGATCCCC |
| Kdm6a F | CCAGAAGTCCTTGGAAGCAGA |
| Kdm6aR | CCGTGGTCCAATTGTACAGC |
| **qPCR primers** | |
| *Gapdh* forward | CTCATGACCACAGTCCATGC |
| *Gapdh* reverse | CACATTGGGGGTAGGAACAC |
| Kdm5c F | GAGGCCCAGACAAGAGTGAAA |
| Kdm5c R | TTGGGAATCTTTAAGGATGAGCC |
| Setd5 F | TCTGTGGAGGCTAGTCCAGC |
| Setd5 R | ATCGTAGCATAAGGCAGTCCT |
| Carm1 F | ATCGCCCTCTACAGCCATGA |
| Carm1 R | CTGTCTGCCCACACGACTG |
